# *In vitro* and *in silico* characterization of human PHOSPHO2 reveals its phospholipid phosphatase activity

**DOI:** 10.64898/2026.07.31.742009

**Authors:** Kyoko Atsuta-Tsunoda, Chiaki Murakami, Hiromichi Sakai, Fumio Sakane

**Author notes:** To whom correspondence should be addressed: C.M., Division of Immune Response, Aichi Cancer Center Research Institute, 1-1 Kanokoden, Chikusa Ward, Nagoya, 464-8681, Japan.; /. These authors contributed equally to this work (Kyoko Atsuta-Tsunoda, Chiaki Murakami).

## Abstract

Phosphatidic acid (PA) phosphatase (PAP) is an enzyme that plays a major role in lipid signaling by controlling the cellular levels of two lipid secondary messengers: its substrate, PA, and its product, diacylglycerol. Two types of mammalian PAPs have been reported to date. Type 1 PAP (PAP1) is an Mg^2+^-dependent, N-ethylmaleimide (NEM)-sensitive cytosolic enzyme (EC 3.1.3.4), whereas type 2 PAP (PAP2), also known as phospholipid phosphatase (PLPP) (EC 3.1.3.113), is an Mg^2+^-independent, NEM-insensitive transmembrane protein. PAP2 also hydrolyzes other bioactive lipids such as lyso-PA (LPA), sphingosine-1-phosphate (S1P), and ceramide-1-phosphate (C1P). Here, we purified human phosphatase orphan 2 (PHOSPHO2), a putative cytosolic phosphatase containing a haloacid dehalogenase-like domain, and characterized its enzymological properties *in vitro*. Purified PHOSPHO2 displays Mg^2+^-dependent, NEM-sensitive phosphatase activities toward PA, LPA, S1P, C1P, and glycerol-3-phosphate (G3P) *in vitro*. Moreover, PHOSPHO2 showed substrate selectivity for PA molecular species containing shorter saturated fatty acids such as lauric acid and myristic acid, or polyunsaturated fatty acids such as docosahexaenoic acid and arachidonic acid. The PAP activity of PHOSPHO2, but not its other phosphatase activities, was strongly enhanced in the presence of phosphatidylcholine and phosphatidylethanolamine, major components of the cell membranes. These results indicate that mammalian PHOSPHO2 is a novel cytosolic PLPP that primarily functions as a PAP on cytoplasm-facing membranes.

## Introduction

Diacylglycerol (DG) is a well-known lipid second messenger that activates protein kinase C (PKC) [1]. DG also regulates a wide variety of signal transduction proteins, including protein kinase D, β2-chimaerin, Munc-13, and Ras guanyl nucleotide-releasing protein [2]. In addition to DG, phosphatidic acid (PA) has been reported to function as a lipid second messenger by regulating various signaling proteins in mammals, including protein kinases such as Raf-1 (C-Raf) kinase, PKCε (novel PKC), PKCζ (atypical PKC), and mammalian target of rapamycin (mTOR), as well as protein phosphatases, lipid kinases, G-protein regulators, and phosphodiesterases [3, 4]. The conversion between DG and PA is therefore a physiologically important reaction.

Phosphatidic acid (PA) phosphatase (PAP) catalyzes the hydrolysis of the linkage between glycerol and phosphate in PA to produce DG and inorganic phosphate [5] (Suppl. Fig. S1 and Fig. 1A). To date, two types of PAPs have been identified in mammals. Type 1 PAP (PAP1) (IUBMB Enzyme Commission Number [6]: EC 3.1.3.4) is an Mg^2+^-dependent and N-ethylmaleimide (NEM)-sensitive cytosolic enzyme, whereas type 2 PAP (PAP2) (phospholipid phosphatase, EC 3.1.3.113) is an Mg^2+^-independent and NEM-insensitive membrane-associated enzyme [7, 8]. PAP1 dephosphorylates PA via a nucleophilic reaction mediated by aspartate residues, with the reaction being facilitated by Mg^2+^ ion [9–11] (Suppl. Fig. S1A). In contrast, PAP2 dephosphorylates PA via charge-relay system involving its catalytic histidine and aspartate residues (Suppl. Fig. S1B) [12]. Therefore, PAP1 and PAP2 catalyze the same reaction, but are regarded as distinct enzymes.

**Figure 1.**
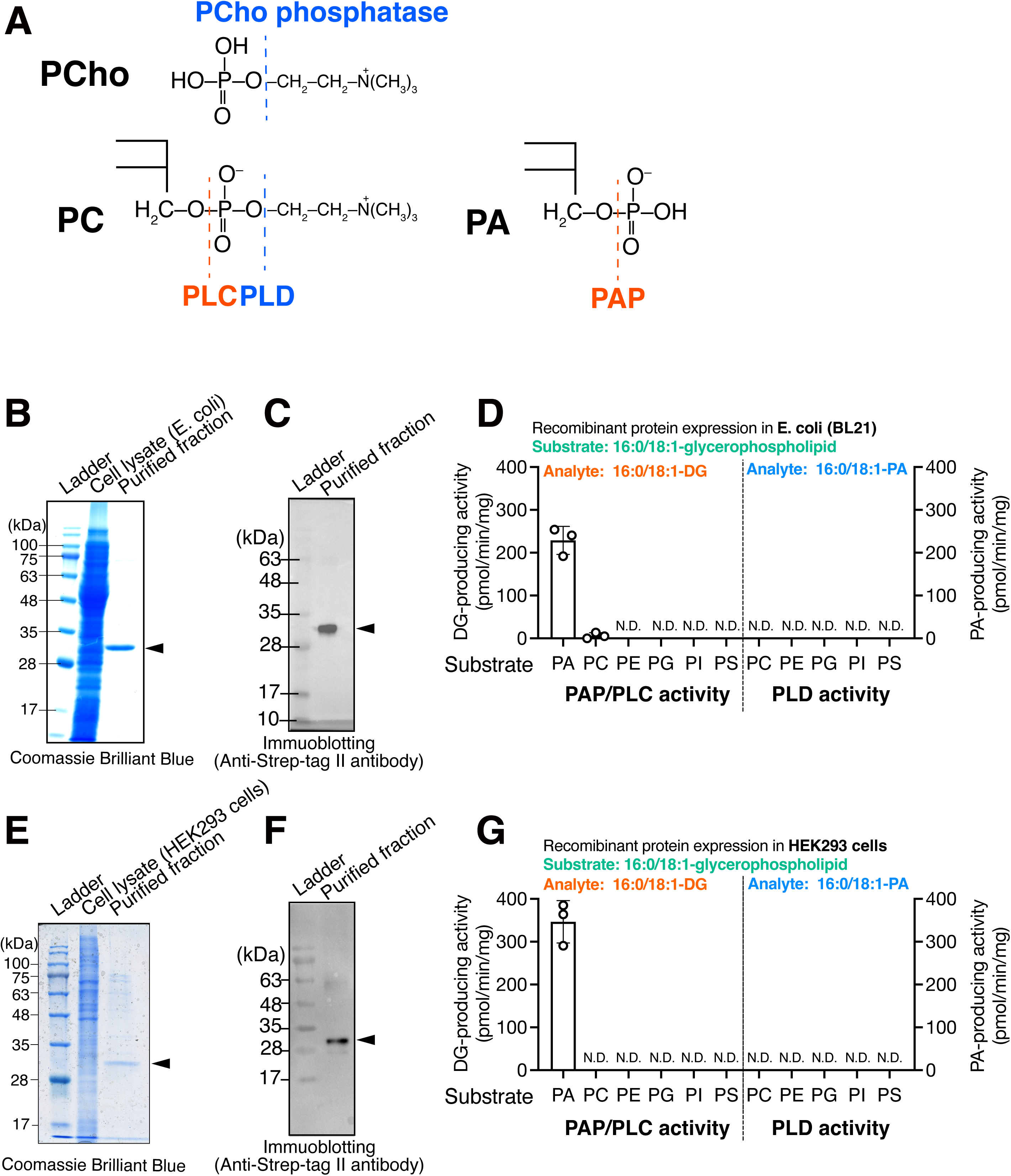
Purification of PHOSPHO2 and in vitro PLC/PLD activity assay. (A) Reactions of phosphatases and hydrolases acting on ester bonds (dotted lines). PHOSPHO1 and PHOSPHO2 display phosphocholine (PCho) phosphatase activity (EC 3.1.3.75) [28]. PC-PLC (EC 3.1.4.3) hydrolyzes PC to produce DG and PCho . Phospholipase D (PLD) (EC 3.1.4.4) hydrolyzes PC to produce phosphatidic acid (PA) and choline (Cho). PAP (EC 3.1.3.4) dephosphorylates (hydrolyzes) PA to produce DG and phosphate. (B–F) C-terminally Twin-Strep-tagged human PHOSPHO2 (PHOSPHO2-TS) was expressed in E.coli BL-21 cells (B, C) or in HEK293 cells (E, F). PHOSPHO2-TS was purified using affinity chromatography with the Strep-Tactin XT beads. Purified PHOSPHO2-TS was analyzed using SDS-PAGE (12% gel), followed by Coomassie Brilliant Blue staining (B, E) [87] or immunoblotting (C, F) using an anti-Strep tag II antibody (1:1,000 dilution in Can Get Signal Solution 1 (Toyobo, Osaka, Japan). BlueStar Prestained Protein Ladder (Cat. No. #NE-MWP03, Nippon Genetics, Tokyo, Japan) was used as a molecular mass marker. DG- and PA-producing activities of PHOSPHO2-TS (500 ng protein/sample) were measured using LC–MS/MS. 16:0/18:1-phospholipids (PC, PE, PA, PI, PG, or PS) in detergent-mixed micelles were incubated with purified PHOSPHO2-TS from BL21 (D) and HEK293 cells (G) for 1 h at 37°C. The peak area ratios of 16:0/18:1-DG (PAP/PLC activities) and 16:0/18:1-PA (PLD activity) in the samples were calculated using an internal standard (I.S.) (0.2 ng/µL of 15:0/18:1-DG and 15:0/18:1-PA). Absolute quantities of 16:0/18:1-DG in samples were determined from calibration curves generated using commercially available purified lipids (16:0/18:1-DG) [41]. Values are presented as means ± S.D. (n = 3, technical replicates). N.D., not detectable.

To date, three mammalian PAP2 genes encoding six-transmembrane proteins have been identified [13, 14]. In addition to PAP activity, all PAP2 enzymes exhibit broad phosphatase activity toward lyso-PA (LPA) (EC 3.1.3.106), sphingosine 1-phosphate (S1P) (EC 3.1.3.114), and ceramide-1-phosphate (C1P) (EC 3.1.3.115) [15]. Because of their broad substrate specificity, PAP2 genes were renamed as phospholipid phosphatases (PLPPs) [16] (Gene names: PLPP1, PLPP2, and PLPP3) (EC 3.1.3.113). Han *et al.* reported that human lipin-1 protein (Gene name: LPIN1) displays Mg^2+^-dependent and NEM-sensitive PAP activity [17, 18]. As PLPPs primarily localize to the plasma membrane, particularly in lipid rafts [19], and their active sites are on the cellular surface [8, 20], PLPPs act as ecto-enzymes that hydrolyze extracellular lipid phosphates. In contrast, lipin proteins are internally localized to the cytosol and nucleus [21]. Both PLPPs and lipins are involved in many signal transduction pathways and a wide variety of biological events, such as glycerolipid biosynthesis, tumorigenesis, obesity, and adipogenesis [5, 8].

Only NEM-insensitive membrane-bound PLPPs were cloned as PAP2. NEM-sensitive C1P phosphatase and LPA phosphatase activities were detected in cytosolic fractions extracted from mammalian tissues [22, 23]. These reports indicate that there are soluble NEM-sensitive PLPP enzymes that are distinct from conventional PLPPs in mammals. However, soluble PLPP genes have not been identified.

The HAD phosphatases are a family of diverse enzymes that are responsible for the majority of metabolic phosphomonoester hydrolysis reactions. Approximately 200 proteins have been identified as HAD phosphatases in *Homo sapiens* [24]. HAD phosphatases employ an aspartate residue in the DXDX(T/V) motif as a nucleophile in a Mg^2+^-dependent phosphoryl transfer reaction (Suppl. Fig. S1A). Although the catalytic motif is conserved in the HAD family, the overall sequence identity between the HAD phosphatases is typically very low (often < 15%), and HAD phosphatases exhibit diverse substrate specificity. Although several HAD phosphatases have been identified and their enzymological properties have been previously characterized, the substrates of some putative HAD phosphatases remain unknown.

Mammalian lipin isoforms (PAP1) are phospholipid phosphatases containing a HAD-like domain, including a DXDXT motif [25]. Phosphatase orphan-1 (Gene name: PHOSPHO1, NCBI Gene ID: 162466, NCBI Reference Sequence: NM_178500.4, UniProt accession number: Q8TCT1) and -2 (Gene name: PHOSPHO2, NCBI Gene ID: 493911, UniProt accession number: Q8TCD6) were initially cloned as putatively soluble cytoplasmic phosphatase enzymes that contain a HAD domain (Suppl. Fig. S2) [26, 27]. Human PHOSPHO1 (267 amino acids, 29.7 kDa) and PHOSPHO2 (241 amino acids, 27.8 kDa) are soluble cytoplasmic proteins that contain no signal peptide [28]. The mRNA of human PHOSPHO1 is predominantly expressed in the testis, whereas that of PHOSPHO2 is ubiquitous [29]. PHOSPHO1 is localized to the plasma membrane and actin filaments, whereas PHOSPHO2 is localized in cellular vesicles [30]. PHOSPHO1 has been implicated in several biological processes, including matrix vesicle-mediated mineralization [31], phospholipid homeostasis [32], and particulate matter-induced energy metabolism disorder [33]. PHOSPHO2 has been linked to the development of hepatocellular carcinoma [34, 35], and tumorigenic processes in gastric cancer [36].

The enzymological properties of PHOSPHO1 and PHOSPHO2 have already been reported. Roberts *et al.* demonstrated that recombinant human PHOSPHO1 and PHOSPHO2 exhibit phosphatase activity toward phosphocholine (PCho)/phosphoethanolamine (PEA) (EC 3.1.3.75) [28] (Fig. 1A) and pyridoxal-5’-phosphate (PLP, the active form of vitamin B_6_) (EC 3.1.3.74) [37], respectively. Although the phosphatase activities of PHOSPHO1 and PHOSPHO2 were measured using only soluble phosphate compounds (*e.g.,* PCho, PEA, pyrophosphate, phosphor-*L*-serine, ATP, and PLP) [28, 37], it remained unclear whether PHOSPHO1 and PHOSPHO2 hydrolyze insoluble phosphorus compounds (*e.g.*, glycerophospholipids such as PA, phosphatidylcholine (PC), and phosphatidylethanolamine (PE)).

Since PHOSPHO1 is a protein that contains HAD domain, catalyzes the hydrolysis of PEA and PCho [37], and is localized to the plasma membrane [30], we hypothesized that PHOSPHO1 possesses phospholipid phosphohydrolase activity and demonstrated that PHOSPHO1 exhibits PC-phospholipase C (PLC) (PC-PLC, EC 3.1.4.3) and PE-PLC (EC 3.1.4.62) activities [38].

The newly discovered enzymological properties of PHOSPHO1 as a PLC [38] led us to hypothesize that PHOSPHO2 may also exhibit PLC activities, including PAP activity (*i.e.* PLC activity toward PA) activity (Fig. 1A). In this study, we purified human PHOSPHO2 and characterized it as a PLC/PAP *in vitro*. PHOSPHO2 displayed Mg^2+^-dependent and NEM-sensitive PAP activity (PAP1 activity). In addition to PA, PHOSPHO2 hydrolyzed phosphate-containing lipids (S1P, C1P and LPA), and glycerol-3-phosphate (G3P) (PAP2/PLPP activity). PHOSPHO2 preferentially hydrolyzed shorter saturated fatty acids (SFAs) such as lauric acid (12:0) (*X*:*Y* = total number of carbon atoms:total number of double bonds) and myristic acid (14:0), or polyunsaturated fatty acid (PUFA)-containing PA, compared to longer SFA-containing PA such as 16:0/16:0-PA and 18:0/18:0-PA.

Collectively, we have revealed the novel enzymological properties of mammalian PHOSPHO2 as a new type of PLPP that is soluble (cytoplasmic enzyme), Mg^2+^-dependent, NEM-sensitive, and selective for shorter SFA-containing or PUFA-containing PA/LPA species.

## Results

### Purification of PHOSPHO2

We first expressed the C-terminally Twin-Strep-tagged human PHOSPHO2 (PHOSPHO2-TS) in both *Escherichia coli* (BL21) (Fig. 1B and C) and mammalian cells (HEK293 cells) (Fig. 1E and F) and purified it from the soluble fractions by affinity chromatography using Strep-Tactin XT [39–41]. A single band with a molecular mass of approximately 32 kDa (calculated PHOSPHO2 mass: 27.8 kDa; TEV site and Twin-Strep-Tag: 4.3 kDa) was observed using Coomassie Brilliant Blue staining (Fig. 1B and E) and immunoblotting with an anti-Strep II tag antibody (Fig. 1C and F). Recombinant PHOSPHO2 was primarily detected in the soluble fraction (Suppl. Fig. S3A). These results indicated that PHOSPHO2-TS with high purity (> 90% purity for the enzyme expressed in *E. coli*) was obtained and that PHOSPHO2 was highly soluble. The final yield of protein was approximately 3.7 μg/L of culture (4.0 × 10^8^ cells in twenty 150-mm dishes) (HEK293 cells) and 200 μg/L of culture (BL21 cells), respectively.

### Enzymological characterization of PHOSPHO2 as a DG-generating enzyme in vitro

We investigated whether PHOSPHO2 exhibits phosphohydrolase activities such as PLC/PAP and phospholipase D (PLD; EC 3.1.4.4) (Fig. 1A) activities toward several phospholipids; 1-palmitoyl-2-oleoyl (16:0/18:1)-PC, -PE, -phosphatidylglycerol (PG), - phosphatidylinositol (PI), -phosphatidylserine (PS), and -PA by measuring the DG-producing activity and PA-producing activity (except for PA), respectively (Fig. 1D and G). Initially, we used LC–MS/MS-based PLC/PAP and PLD activity assays [38, 39, 41–43]. All enzyme activity assays were performed in detergents (n-dodecyl-β-D-maltoside (DDM) and cholesteryl hemisuccinate (CHS)-based micelle environment.

Recombinant human PHOSPHO2 (500 ng protein/sample), which was expressed in and purified from both *E. coli* (Fig. 1D) and mammalian (HEK293) cells (Fig. 1G), produced DG in the presence of PA-containing micelles. These results indicated that, *in vitro,* human PHOSPHO2 possesses PAP (PA-PLC) activity but not PLD activity. Moreover, this enzyme failed to show PC-, PE-, PG-, PI-, and PS-PLC activities (Fig. 1D and G). The enzymological properties (*e.g.,* substrate selectivity and specific activity) of PHOSPHO2 expressed in *E. coli* (Fig. 1D) are similar to those of the protein expressed in mammalian cells (Fig. 1G). Moreover, the yield of PHOSPHO2-TS expressed in *E. coli* was higher than that of mammalian cells (200 μg/L versus 3.7 μg/L). Because PHOSPHO2 displayed PAP activity, we employed a phosphatase activity assay using BIOMOL Green for subsequent experiments. First, we confirmed that the PA phosphatase reaction catalyzed by PHOSPHO2 is linear with time for at least 120 min (Suppl. Fig. S3B), indicating that the reaction followed zero-order kinetics with respect to PA within the DDM/CHS-mixed micelles. However, the colorimetric method had lower sensitivity than the LC-MS/MS method, necessitating the use of a larger amount of recombinant protein. Therefore, routine phosphatase activity assays were performed for 60 min using 4 µg of recombinant protein purified from *E. coli*.

### Fatty acyl chain selectivity of PHOSPHO2 toward PA molecular species

The effects of acyl chains of PA on the PAP activity of PHOSPHO2 were examined (Fig. 2). Purified PHOSPHO2 exhibited high PAP activity toward shorter SFA- or PUFA-containing PA molecular species such as 12:0/12:0-, 14:0/14:0-, 18:0/20:4-, and 18:0/22:6-PA. PAP activity toward 12:0/12:0-PA ranged 3–8-fold greater than the activities toward longer SFA (16:0 or 18:0)- and/or monounsaturated fatty acid (MUFA) (18:1)-containing PA molecular species such as 16:0/16:0-, 18:0/18:0-, 16:0/18:1-, and 18:1/18:1-PA.

**Figure 2.**
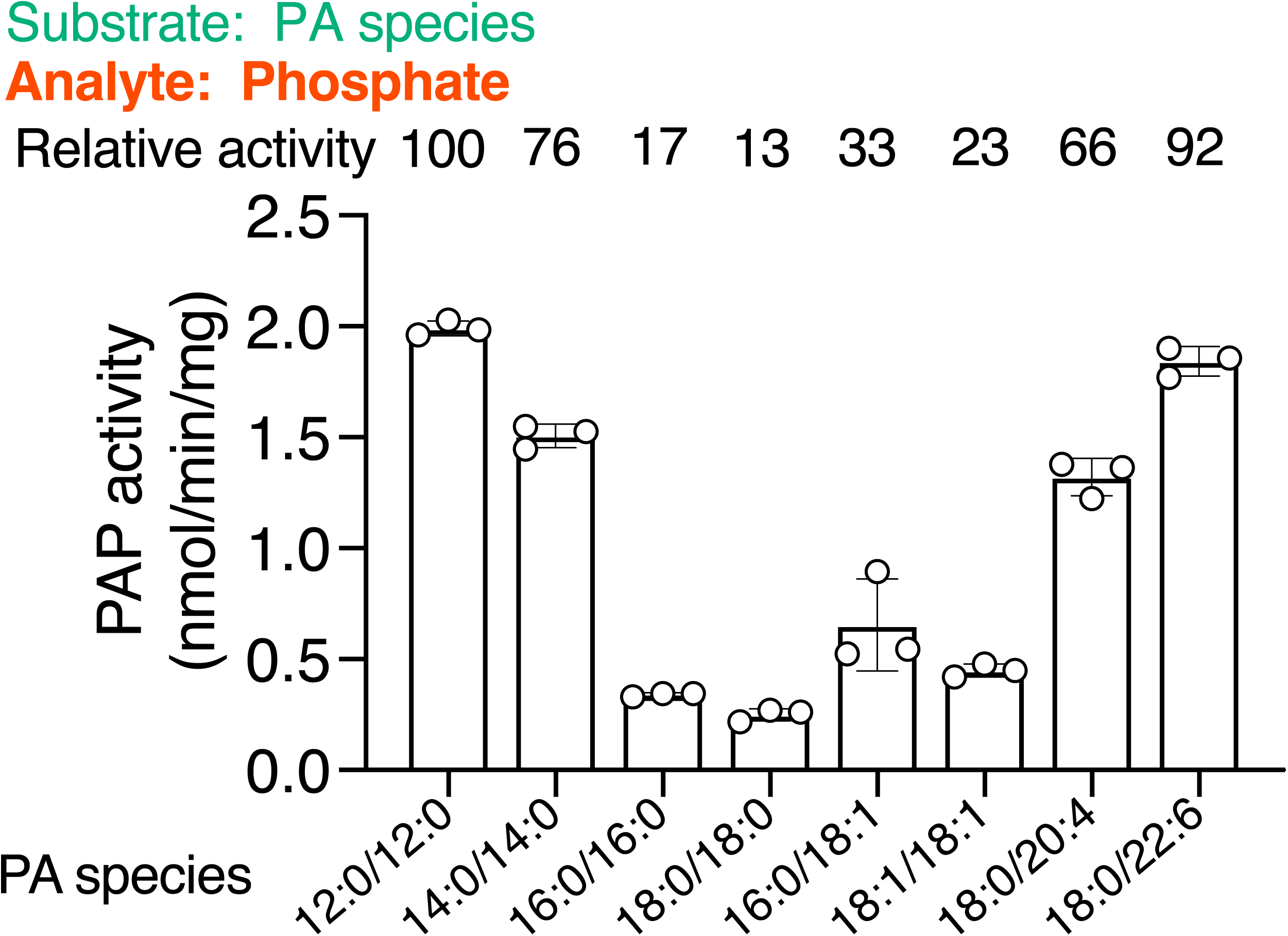
Effects of fatty acyl chains of PA on the PAP activity of PHOSPHO2. Comparison of PAP activities of PHOSPHO2-TS toward various PA molecular species (12:0/12:0-, 14:0/14:0-, 16:0/16:0-, 18:0/18:0-, 16:0/18:1-, 18:1/18:1-, 18:0/20:4-, and 18:0/22:6-PA). To compare PAP activities toward various PA species, free inorganic phosphate released from the substrates was measured using the BIOMOL Green reagent. Values are presented as means ± S.D. (*n* = 4, technical replicates).

### Effects of temperature and pH on the PAP activity of PHOSPHO2

The effect of temperature on the PAP activity of PHOSPHO2 was examined (Fig. 3A). PHOSPHO2 exhibited PAP activity in the range of 30–50°C and was essentially inactive at 10, 20, and 60°C. Maximum activity was observed at 40°C. The thermal stability of PHOSPHO2 was examined by preincubation for 20 min at 0, 30, 40, 50, and 60°C (Fig. 3B). Following the preincubation, the purified PHOSPHO2 samples were cooled on ice for 10 min, and then assayed for PAP activity. More than 50% of the PAP activity of PHOSPHO2 was retained after incubation at temperatures below 50°C. PAP activity was completely halted by preincubation at 60°C.

**Figure 3.**
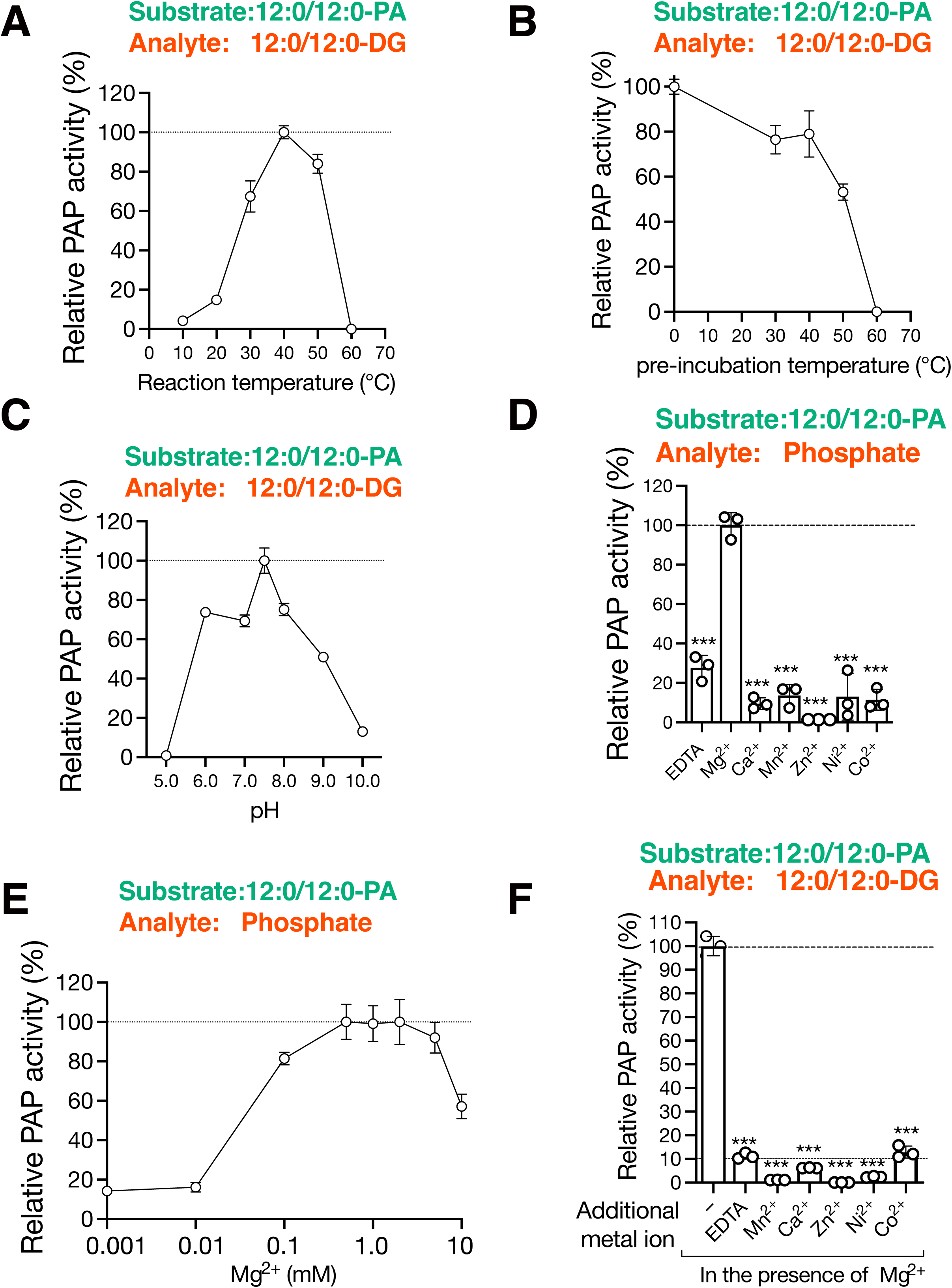
Effects of temperature, pH, and divalent metal ions on PAP activity and stabilities of PHOSPHO2. (*A*) Using purified PHOSPHO2-TS expressed in *E. coli* BL21, PAP activity was measured at the indicated temperatures (10, 20, 30, 40, 50, and 60°C) for 2 h using 12:0/12:0-PA at a final concentration of 200 µM. Relative activities are presented as percentages of the PAP activities at 40°C (set to 100%). Values are presented as means ± S.D. (*n* = 3, technical replicates). *(B)* PHOSPHO2-TS was pre-incubated at the indicated temperatures (0, 30, 40, 50, and 60°C) for 20 min. After preincubation, the samples were cooled in an ice bath for 5 min to allow for enzyme renaturation, and PAP activity was then measured for 2 h at 37°C. The highest activity preincubated at 0°C was set to 100%. Values are presented as means ± S.D. (*n* = 3, technical replicates). *(C)* PAP activity was measured at the indicated pH values (pH 5.0, 6.0, 7.0, 7.5, 8.0, 9.0, and 10.0) with 50 mM Tris-maleate-NaOH buffer. Values are presented as the mean ± S.D. (*n* = 3, technical replicates). *(D)* Effects of divalent metal ions (Mg^2+^, Ca^2+^, Mn^2+^, Zn^2+^, Ni^2+^, Co^2+^) or EDTA (each at 1 mM) on the PAP activity of PHOSPHO2 purified in the absence of Mg^2+^ were determined using the LC–MS/MS-based enzyme activity assay [38, 41, 42]. Relative activities are presented as percentages of PAP activities with 1 mM of Mg^2+^ (set to 100%). Values are presented as means ± S.D. (*n* = 3, technical replicates). ***, *p* < 0.001 (*vs.* Mg^2+^). One-way ANOVA with Dunnett’s post hoc test was used. *(E)* PAP activity was measured in the presence of indicated concentrations of MgCl_2_ (0.001, 0.01, 0.1, 0.5, 1, 2, 5, and 10 mM). Relative activities are presented as percentages of PAP activities with 1 mM of Mg^2+^ (set to 100%). Values are presented as means ± S.D. (*n* = 3, technical replicates). *(F)* In the presence of Mg^2+^ (1 mM), PAP activity was measured with 1 mM of Ca^2+^, Mn^2+^, Zn^2+^, Ni^2+^, Co^2+^, or EDTA. PHOSPHO2-TS was expressed in *E. coli* (BL21) and purified. Values are presented as means ± S.D. (*n* = 3, technical replicates). Relative activities are presented as percentages of the PAP activities with 1 mM of Mg^2+^ (set to 100%). ***, *p* < 0.005 (*vs.* control).

The effect of pH on the PAP activity of PHOSPHO2 was determined (Fig. 3C). PHOSPHO2 exhibited PAP activity in the range of pH 6.0–9.0, with optimum activity at pH 7.5. The pH optimum for PHOSPHO2 (pH 7.5) was higher than that for PHOSPHO1 (around pH 6.7) [37].

### Effects of divalent metal ions on PAP activity of PHOSPHO2

Because the DxDxT motif, a Mg^2+^-binding domain [9], is conserved in PHOSPHO2 (Suppl. Fig. S2), we tested the effects of divalent metal ions (1 mM of Mg^2+^, Ca^2+^, Mn^2+^, Zn^2+^, Ni^2+^, and Co^2+^) on PAP activity (Fig. 3D). PHOSPHO2 was purified in the absence of MgCl_2_ and the effect of divalent metal ions on PAP activity of PHOSPHO2 was evaluated by incubation of 16:0/18:1-PA-containing micelles with 1 mM concentrations of various metal ions. As a control, PHOSPHO2 was incubated with 16:0/18:1-PA containing 1 mM EDTA. As shown in Fig. 3D, PAP activity was highest in the presence of 1 mM Mg^2+^. Compared with the metal ion-free control, Ca^2+^, Mn^2+^, Ni^2+^, and Co^2+^ did not enhance PAP activity, whereas the presence of Zn^2+^ strongly inhibited the PAP activity (the order of activity enhancement: Mg^2+^ >> no metal ≈ Mn^2+^ ≈ Ni^2+^ ≈ Ca^2+^ ≈ Co^2+^ >> Zn^2+^).

The optimum Mg^2+^ concentration of the PAP activity was determined (Fig. 3E). PHOSPHO2 exhibited strong PAP activity in the range of 0.1–10 mM MgCl_2_, with optimum activity at 0.5–2 mM MgCl_2_. At high Mg^2+^ concentrations (10 mM), PAP activity instead decreased.

Zn^2+^ reportedly inhibits Lipin-1 [17], and mammalian PLPPs are inhibited by divalent metal ions such as Zn^2+^, Mn^2+^, Ca^2+^, and Co^2+^[7, 44]. This led us to evaluate the effects of several metal ions other than Mg^2+^ on PAP activity in the presence of 1 mM Mg^2+^ (Fig. 3F). The activity was also inhibited (90–98% inhibition, but not 100%) by the addition of 1 mM Ca^2+^, Mn^2+^, Ni^2+^, Co^2+^, Zn^2+^ or EDTA.

### Kinetic properties of PHOSPHO2 as PAP

The kinetic analyses of PHOSPHO2 were conducted. First, we determined the kinetic constant based on PA molar concentration using DDM/CHS/PA-mixed micelles (Fig. 4). In this experiment, bulk 12:0/12:0-PA concentration was varied (0.025–0.4 mM) by maintaining the molar ratio of detergents to PA in the micelles at 9:1 (*i.e.* 10 mol% PA). The *K*_m_ and *V*_max_ were calculated to 798 ± 102 μM and 10.5 ± 1.0 nmol/min/mg, respectively, and are broadly comparable with those of Lipin-1s [17] (Table 1).

**Figure 4.**
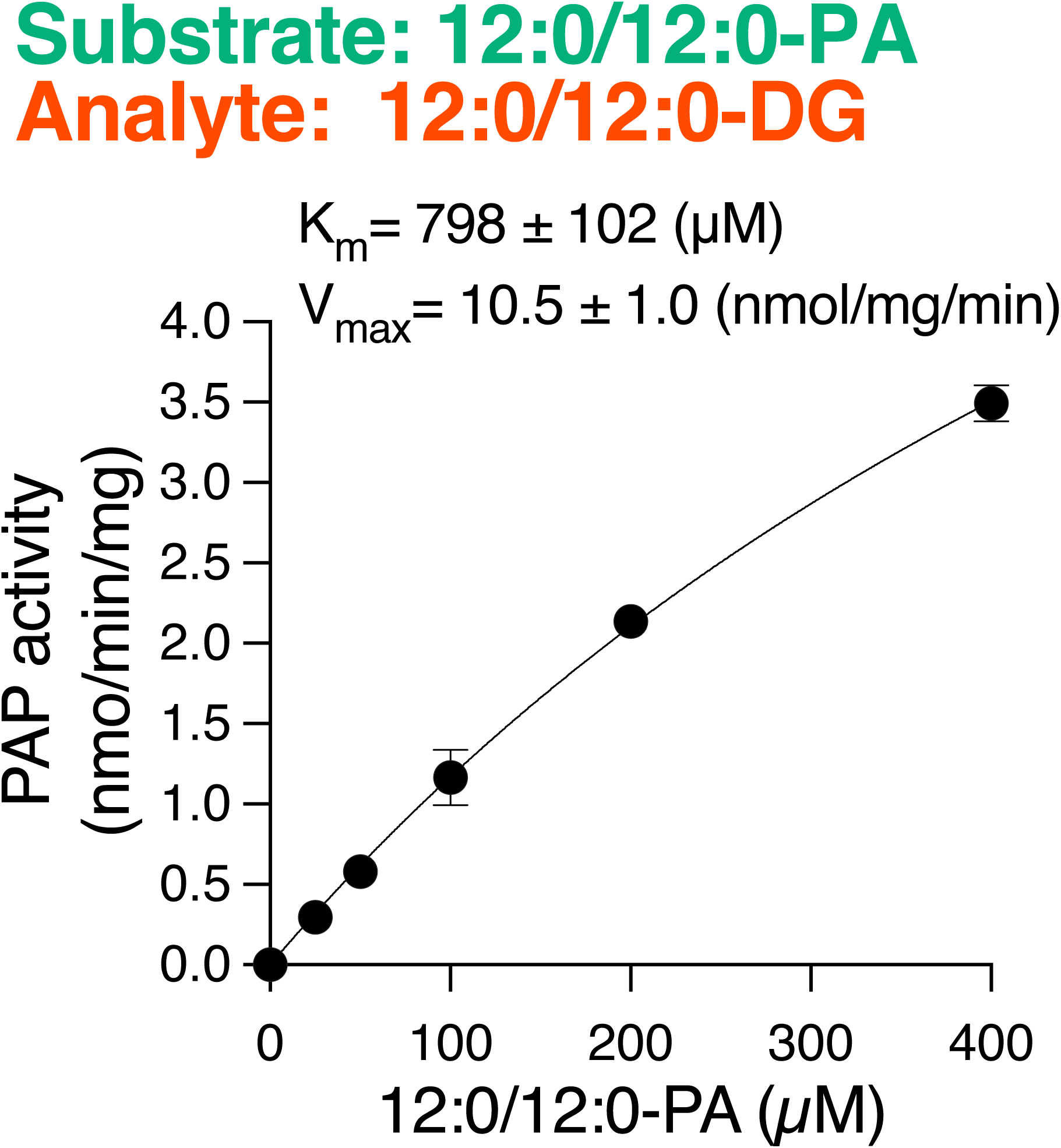
Kinetic analysis of PAP activities of PHOSPHO2. PAP activity was measured as a function of the indicated bulk molar concentration (µM) using 12:0/12:0-PA-DDM/CHS mixed micelles. In the experiment, 12:0/12:0-PA bulk concentration was varied (0.025–0.4 mM). The PAP (DG-producing) activities of PHOSPHO2 were measured using LC-MS/MS [38, 41, 42]. Data are presented as means ± S.D. (*n* = 3, technical replicates). The enzyme kinetic parameters (*K*_m_ and *V*_max_) were obtained by fitting kinetic models to the experimental data using GraphPad Prism 9.

**Table 1.** Comparison of PHOSPHO2 (our findings and previous reports), Lipin, and PLPPs.

| Properties | PHOSPHO2<br>(Our findings) | Phosphatase orphan<br>(previous reports) | Type 1 PAP<br>(Mg<sup>2+</sup>-dependent soluble<br>PAP) | Type 2 PAP<br>(Mg<sup>2+</sup>-<br>independent<br>membrane<br>associated PAP) |
| --- | --- | --- | --- | --- |
| Identified<br>Genes |  | PHOSPHO1 [37]<br>PHOPSHO2 [28] | LPIN1 [17]<br>LPIN2 [18]<br>LPIN3 [18] | LPP1/PAP-2a<br>(PLPP1) [13]<br>LPP2/PAP-2c<br>(PLPP2) [14]<br>LPP3/PAP-2b<br>(PLPP3) [13] |
| Subcellular<br>localization | Cytosolic fraction | **PHOSPHO1 :**<br>Cytosolic fraction [28, 37]<br>plasma membrane [30]<br>actin filaments, [30]<br>**PHOSPHO2:**<br>Cytosolic fraction [28]<br>vesicles [30] | Cytosol [92]<br>ER membrane [92] | Plasma<br>membrane (raft)<br>[19] |
| Mg<sup>2+</sup><br>dependency | Dependent | Dependent [28, 37]: | Dependent [17] | Independent [7] |
| Enzyme<br>reaction | PA + H<sub>2</sub>O→DG + Pi<br>LPA + H<sub>2</sub>O→MG + Pi<br>S1P + H<sub>2</sub>O→Sph + Pi<br>C1P + H<sub>2</sub>O→Cer + Pi<br>GP + H<sub>2</sub>O→Glycerol+ Pi<br>PLP + H<sub>2</sub>O→Pyridoxal + Pi | **PHOSPHO1** [37, 38]:<br>PEA + H<sub>2</sub>O→Ethanolamine<br>+ Pi<br>PCho + H<sub>2</sub>O→Choline + Pi<br>PC + H<sub>2</sub>O→DG + PCho<br>PE + H<sub>2</sub>O→DG + PEA<br>**PHOSPHO2**[28]: | PA + H<sub>2</sub>O→DG + Pi | PA + H<sub>2</sub>O→DG + Pi<br>LPA + H<sub>2</sub>O→MG +<br>Pi<br>S1P + H<sub>2</sub>O→Sph +<br>Pi<br>C1P + H<sub>2</sub>O→Cer +<br>Pi |
|  |  | PLP + H<sub>2</sub>O→Pyridoxal + Pi |  | DGPP + H<sub>2</sub>O→PA + Pi<br>[13, 15] |
| Relative activity | d18:1-S1P: 100<br>PLP: 91<br>12:0/12:0-PA: 78<br>G3P: 26<br>16:0-LPA: 15<br>d18:1/18:0-C1P:13<br>(Fig. 6A) | **PHOSPHO1**[28, 37, 38]:<br>PEA: 100<br>PCho: 65<br>PC: 0.2<br>PE: 0.2<br>**PHOSPHO2** [28, 37]:<br>PLP: 100<br>ATP: 30<br>Phosphor-L-serine: 30 | PA: 100<br>(hydrolyze only PA) [17] | **LPP1/PAP-2a** [7]:<br>PA:100<br>LPA: 93<br>S1P: 59<br>C1P: 56<br>**LPP2/PAP-2c** [7]:<br>S1P: 100<br>LPA: 29<br>C1P: 25<br>PA: 22<br>**LPP3/PAP-2b** [7]:<br>LPA: 100<br>C1P: 78<br>PA: 59<br>S1P: 52 |
| Kinetic parameter (*K<sub>m</sub>*) | PLP: > 3,000<br>12:0/12:0-PA: 798 ± 102 μM (3.06 ± 0.89 mol%)<br>16:0/18:1-PA: 30.9 ± 6.0 μM<br>24:0-LPA: 624 ± 126 μM<br>d18:1/8:0-C1P: 117 ± 42 μM | **PHOSPHO1**<br>(Roberts *et al.*) [37]<br>PEA: 3.0 μM<br>PCho: 11.4 μM<br>(Murakami *al.*) [38] | **LPIN1 isoforms** [17]:<br>**Lipin-1α**<br>PA: 350 μM (4.2 mol%)<br>**Lipin-1β:**<br>PA: 240 μM (4.5 mol%) | **LPP1/PAP-2a** [7]:<br>PA: 3.4 mol%<br>LPA: 1.3 mol%<br>S1P: 2.8 mol%<br>C1P: 7.1 mol% |
| | d16:1-S1P: $160 \pm 20 \mu\text{M}$<br>d17:1-S1P: $27 \pm 5 \mu\text{M}$<br>d18:1-S1P: $73 \pm 22 \mu\text{M}$<br>d20:1-S1P: $23 \pm 14 \mu\text{M}$<br>(Fig. 4 and 11) | PC > $400 \mu\text{M}$<br>PCho > $1,000 \mu\text{M}$<br>**PHOSPHO2** [28]:<br>PLP: $45.5 \mu\text{M}$ | **Lipin-1γ**:<br>PA: $110 \mu\text{M}$ (4.3 mol%) | **LPP2/PAP-2c** [7]:<br>S1P: 5.99 mol%<br>LPA: 1.35 mol%<br>C1P: 0.78 mol%<br>PA: 0.44 mol%<br>**LPP3/PAP-2b** [7]:<br>LPA: 1.11 mol%<br>C1P: 3.4 mol%<br>PA: 0.61 mol%<br>S1P: 2.5 mol% |
| Maximum rate of reaction ($V_{\text{max}}$) in mixed-micelle activity assay (nmol/min/mg) | 16:0/18:1-PA: $1.0 \pm 0.05$<br>12:0/12:0-PA: $10.5 \pm 1.0$<br>24:0-LPA: $64.9 \pm 8.9$<br>PLP: > 40<br>d18:1/8:0-C1P: $15.8 \pm 2.3$<br>d16:1-S1P: $85.2 \pm 4.8$<br>d17:1-S1P: $53.3 \pm 2.3$<br>d18:1-S1P: $15.7 \pm 1.5$<br>d20:1-S1P: $11.4 \pm 1.4$<br>(Fig. 4 and 11) | **PHOSPHO1**:<br>(Roberts *et al.*) [37]<br>PEA: 4,120<br>PCho: 3,600<br>(Murakami *al.*) [38]<br>PC : 1.47<br>PCho : 570<br>**PHOSPHO2**[28]:<br>PLP: 633 | **LPIN1 isoforms** [17]:<br>**Lipin-1α**:<br>PA: > 30<br>**Lipin-1β**:<br>PA: > 20<br>**Lipin-1γ**:<br>PA: > 3 | **LPP1/PAP-2a** [7]:<br>PA: 0.54<br>LPA: 0.50<br>S1P: 0.32<br>C1P: 0.30<br>**LPP2/PAP-2c** [7]:<br>S1P: 0.69<br>LPA: 0.20<br>C1P: 0.17<br>PA: 0.15<br>**LPP3/PAP-2b** [7]:<br>LPA: 0.46<br>C1P: 0.36<br>PA: 0.27<br>S1P: 0.24 |
| Acyl chain specificity (*in vitro* study) | PUFA (20:4) > shorter SFA (12:0/12:0 and 14:0/14:0) > | Unknown | Unsaturated FA-containing PA (18:1, 18:2, 20:4 or 22:6-containing PA) >>> SFA-PA | Unknown |
|  | MUFA (18:1 containing) > SFA (16:0/16:0, 18:0/18:0) (Fig. 2) |  | (16:0/16:0- and 18:0/18:0-PA) [17] |  |
| Propranolol sensitivity | Partly Sensitive (PA and C1P) (Fig. 5A and 9) | Unknown | Sensitive [17, 93, 94] | Sensitive [7, 13]<br>**IC<sub>50</sub> [7]:**<br>**LPP1/PAP-2a:**<br>≈3 mM<br>**LPP2/PAP-2c:**<br>≈1 mM<br>**LPP3/PAP-2b:**<br>≈2 mM |
| NEM sensitivity | Sensitive (PLP, PA, LPA, C1P and S1P) (Fig. 5A and 9) | Unknown | Sensitive [17, 95] | Insensitive [7, 95] |
| Other inhibitors | Zn<sup>2+</sup>, Mn<sup>2+</sup>, Ca<sup>2+</sup>, Co<sup>2+</sup>, Ni<sup>2+</sup>, EDTA, Sph, PI, Cer (Fig. 3F and 5B) | D609 (inhibited PLC activity of PHOSPHO1) [38] | Na<sub>3</sub>VO<sub>4</sub>, CaCl<sub>2</sub>, ZnCl<sub>2</sub>, MnCl<sub>2</sub>, Sph, sphinganine, C1P [17, 94, 95] | Zn<sup>2+</sup>, Sph [7, 94, 95] |

The PA phosphatase activity assay using micelles permitted the analysis of kinetic properties in an environment that mimicked the surface of the cellular membrane (also known as “surface dilution kinetics model”) [45]. Therefore, we determined the *K*_m_ and *V*_max_ of PHOSPHO2 for PA using the surface dilution kinetics model (Suppl. Fig. S4). In this experiment, PAP activity of PHOSPHO2 was measured as a function of the surface concentration of PA in the micelles (mol% = 100 × [PA (mol)]/([DDM (mol)] + [CHS (mol)] + [PA (mol)])) by maintaining the molar concentration of 12:0/12:0-PA at 0.2 mM. Under the conditions of the assay, PAP activity was dependent on the surface concentration of PA (mol%) but independent of the bulk PA concentration (mM), suggesting that the PAP activity of PHOSPHO2 follow the surface dilution kinetic model.

### Effects of PAP inhibitors on the PAP activity of PHOSPHO2

NEM inhibits only PAP1/Lipin-1 [7, 18]. Propranolol, a non-selective β-blocker, is an inhibitor for both PAP1/Lipin-1 and PAP2/PLPP [7, 17] (Table 1). Therefore, the effects of NEM and propranolol on the PAP activity of PHOSPHO2 were examined (Fig. 5A). Both NEM and propranolol inhibited the PAP activity of PHOSPHO2 in a dose-dependent manner. The IC_50_ values of NEM and Propranolol were 0.26 mM and 0.30 mM, respectively, indicating that PHOSPHO2 is sensitive to both NEM and propranolol.

**Figure 5.**
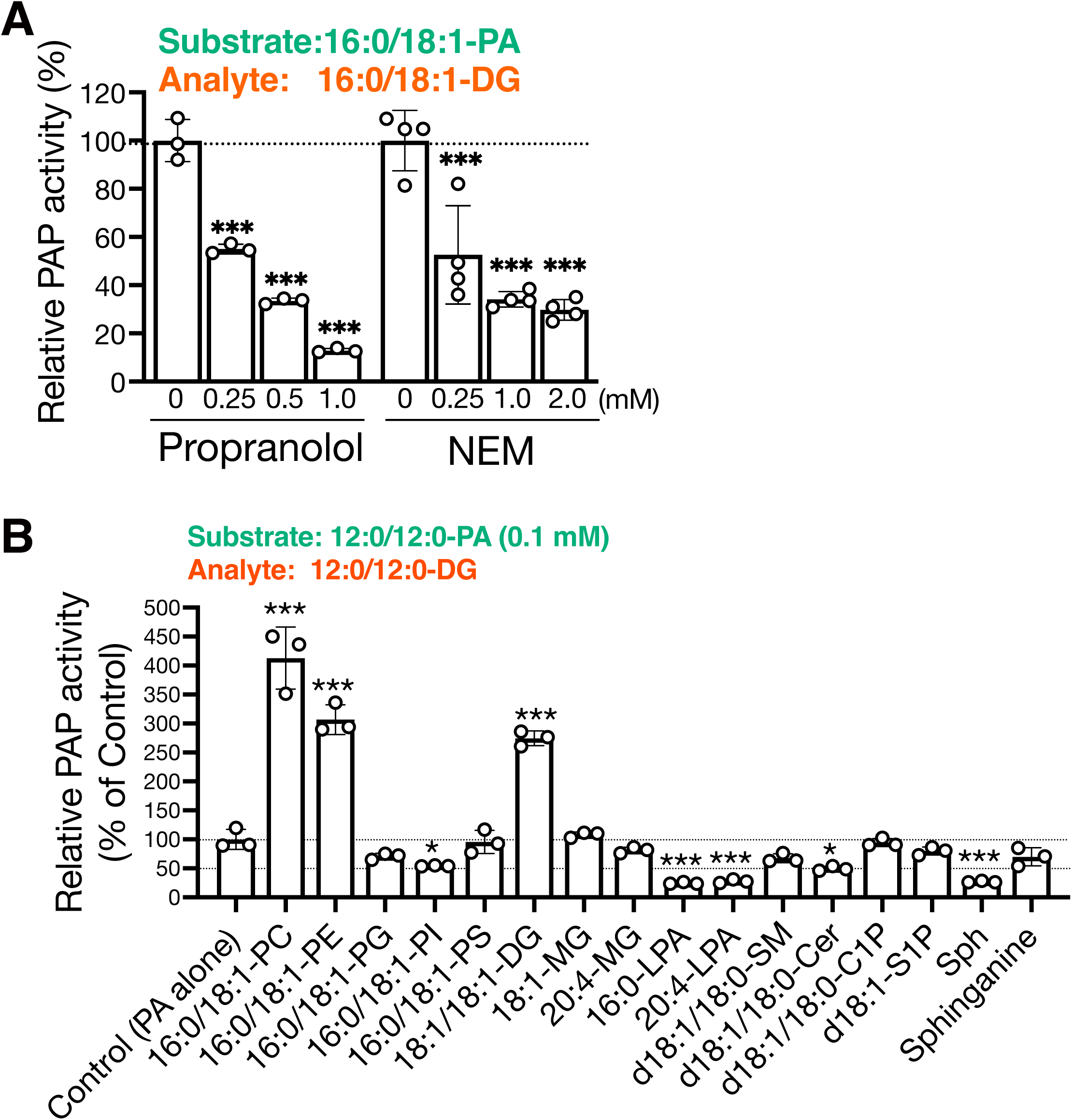
Effects of PAP inhibitors and various lipids on PAP activities of PHOSPHO2. (*A*) PAP activity of PHOSPHO2 was measured in the presence of various concentrations of *N*-ethylmaleimide (NEM) or propranolol. Purified PHOSPHO2-TS (approximately 500 ng) was incubated with 12:0/12:0-PA-detergents mixed micelles for 1 h at 37°C. Produced 12:0/12:0-DG was quantified using LC–MS/MS. Values are presented as percentages of the activity of PHOSPHO2 in the absence of inhibitor (set at 100%). Values are presented as means ± S.D. (*n* = 3–4, technical replicates). ***, *p* < 0.005 (*vs.* control (without inhibitor)). One-way ANOVA with Dunnett’s post hoc test was used. (*B*) PAP activity of PHOSPHO2 was measured in the presence of various lipids (each at 0.1 mM) in addition to 12:0/12:0-PA (0.1 mM) as the substrate. The produced 12:0/12:0-DG was quantified using LC–MS/MS. Values are presented as percentages of PAP activity of PHOSPHO2 in the presence of PA-mixed micelle alone (control) (set at 100%). Values are presented as means ± S.D. (*n* = 3, technical replicates). *, *p* < 0.05; ***, *p* < 0.005 (*vs.* control (without inhibitor)). One-way ANOVA with Dunnett’s post hoc test was used.

### Effects of various lipids on the PAP activity of PHOSPHO2

Sphingosine (Sph) inhibits PAP activity of PAP1/Lipin-1 and PAP2/PLPP [7, 17]. Therefore, we determined whether various lipids, including Sph, affect the PAP activity of PHOSPHO2 (Fig. 5B). Sph and LPA strongly inhibited PAP activity (causing a decrease of approximately 73% and 75%, respectively). Ceramide (Cer) and PI significantly attenuated PAP activity (approximately 50% and 46% decrease, respectively). DG, PC, and PE strongly enhanced the PAP activity (approximately 2.5-, 4.0-, and 3.0-fold, respectively). However, PS, PG, sphingomyelin (SM), ceramide 1-phosphate (C1P), sphingosine 1-phosphate (S1P), monoacylglycerol (MG), and sphinganine did not affect PAP activity.

### Substrate specificity of PHOSPHO2 toward phosphate-containing lipids

We then tested whether PHOSPHO2 hydrolyzes (dephosphorylates) various phosphorus compounds (LPA, S1P, C1P, and G3P) in addition to PA and PLP. As shown in Fig. 6A, d18:1-S1P was dephosphorylated by PHOSPHO2 with the highest specific activity, which was comparable to the phosphatase activities toward 12:0/12:0-PA and PLP. PHOSPHO2 also hydrolyzed G3P, 16:0-LPA, and d18:1/18:0-C1P (approximately 30%, 17%, and 14% of PLP phosphatase activity). These results suggest that PHOSPHO2 acts as a cytosolic PLPP.

**Figure 6.**
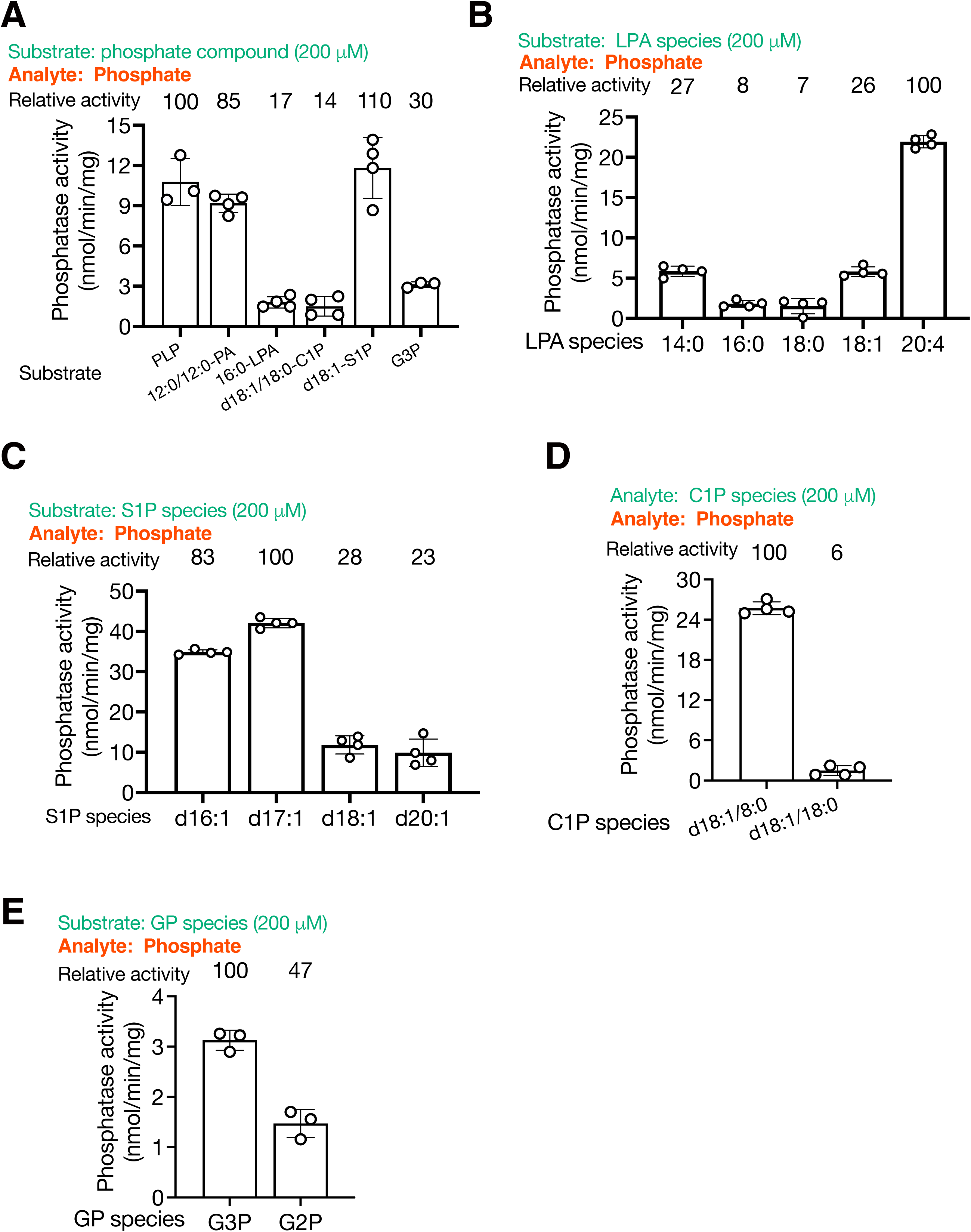
Specific phosphatase activities of PHOSPHO2 toward phosphorus compounds. (A) PHOSPHO2-TS was incubated with a range of substrates (0.2 mM of PLP, 12:0/12:0-PA, 16:0-LPA, d18:1/18:0-C1P, d18:1-S1P, or G3P). To compare phosphatase activities, free inorganic phosphate released from the substrates was measured using the BIOMOL Green reagent. (B–E) Comparison of phosphatase activities toward LPA molecular species (14:0-, 16:0-, 18:0-, 18:1-, and 20:4-LPA) (B), S1P molecular species (d16:1-, d17:1-, d18:1-, and d20:1-S1P) (C). C1P molecular species (d18:1/8:0-, and d18:1/18:0-C1P) (D), and glycerophosphates (GPs) (G3P and G2P) (E). Values are presented as means ± S.D. (*n* = 3– 4, technical replicates).

### Fatty acyl chain specificities of PHOSPHO2 toward LPA, S1P, and C1P

The effects of acyl chains of glycerophospholipids on phosphatase activities of PHOSPHO2 were examined (Fig. 6B–D). As shown in Fig. 6B, PHOSPHO2 displayed higher phosphatase activity toward PUFA- or shorter SFA-containing LPA molecular species (the order of LPA-phosphatase activity: 20:4 >> 14:0 ≈ 18:1 > 16:0 ≈ 18:0). More specifically, PHOSPHO2 displayed the highest phosphatase activity toward PUFA-containing LPA (20:4-LPA), approximately 3-fold higher than for 14:0- and 18:1-LPA. Among SFA-containing LPA species (14:0-, 16:0-, and 18:0-LPA), PHOSPHO2 exhibited higher phosphatase activity toward shorter SFA-containing LPA (14:0-LPA). As shown in Fig. 6C, we observed that PHOSPHO2 preferably hydrolyzed shorter sphingoid base-containing S1P (the order of S1P-phosphatase activity: d16:1 ≈ d17:1 > d18:1 ≈ d20:1). PHOSPHO2 exhibited higher phosphatase activity toward shorter fatty acid chain-containing d18:1-C1P (the order of C1P-phosphatase activity: d18:1/8:0 > d18:1/18:0) (Fig. 6D). We compared the phosphatase activities of PHOSPHO2 toward GP molecular species, G3P (α-glycerophosphate, a major isomer) and G2P (β-glycerophosphate, less common isomer) (Fig. 6E). PHOSPHO2 hydrolyzed both isomers. However, the phosphatase activity toward G3P was 2-fold higher than that of G2P.

### Kinetic analysis of phosphatase activities of PHOSPHO2 against phosphate compounds presented in detergent mixed micelles

The Michaelis–Menten constant (*K*_m_) and *V*_max_ of PHOSPHO2 as a phosphatase for lipid phosphates (24:0-LPA, d18:1/8:0-C1P, d16:1-S1P, d17:1-S1P, d20:1-S1P, and 16:0/18:1-PA), PLP, and G3P were determined (Table2 and Fig. 7). In this assay, DDM/CHS-mixed micelles were included to maintain the same experimental conditions as those used in the PAP activity assay (Fig. 4). Phosphatase activities increased in a substrate-dependent manner. The *K*_m_ and *V*_max_ against the substrates were as follows: 24:0-LPA, 624 ± 126 μM and 64.9 ± 8.9 nmol/min/mg; d18:1/8:0-C1P, 117 ± 42 μM and 15.8 ± 2.3 nmol/min/mg; d16:1-S1P, 160 ± 20 μM and 85.2 ± 4.8 nmol/min/mg; d17:1-S1P, 27 ± 5 μM and 53.3 ± 2.3 nmol/min/mg; d18:1-S1P, 73 ± 22 μM and 15.7 ± 1.5 nmol/min/mg; d20:1-S1P, 23 ± 14 μM and 11.4 ± 1.4 nmol/min/mg; 16:0/18:1-PA, 30.9 ± 6.0 μM and 1.0 ± 0.05 nmol/min/mg; G3P, 2.2 ± 0.3 mM and 8.2 ± 0.6 nmol/min/mg, respectively. However, we could not determine *K*_m_ (>3,000 μM) or *V*_max_ (> 40 nmol/min/mg protein) of PHOSPHO2 toward PLP.

**Figure 7.**
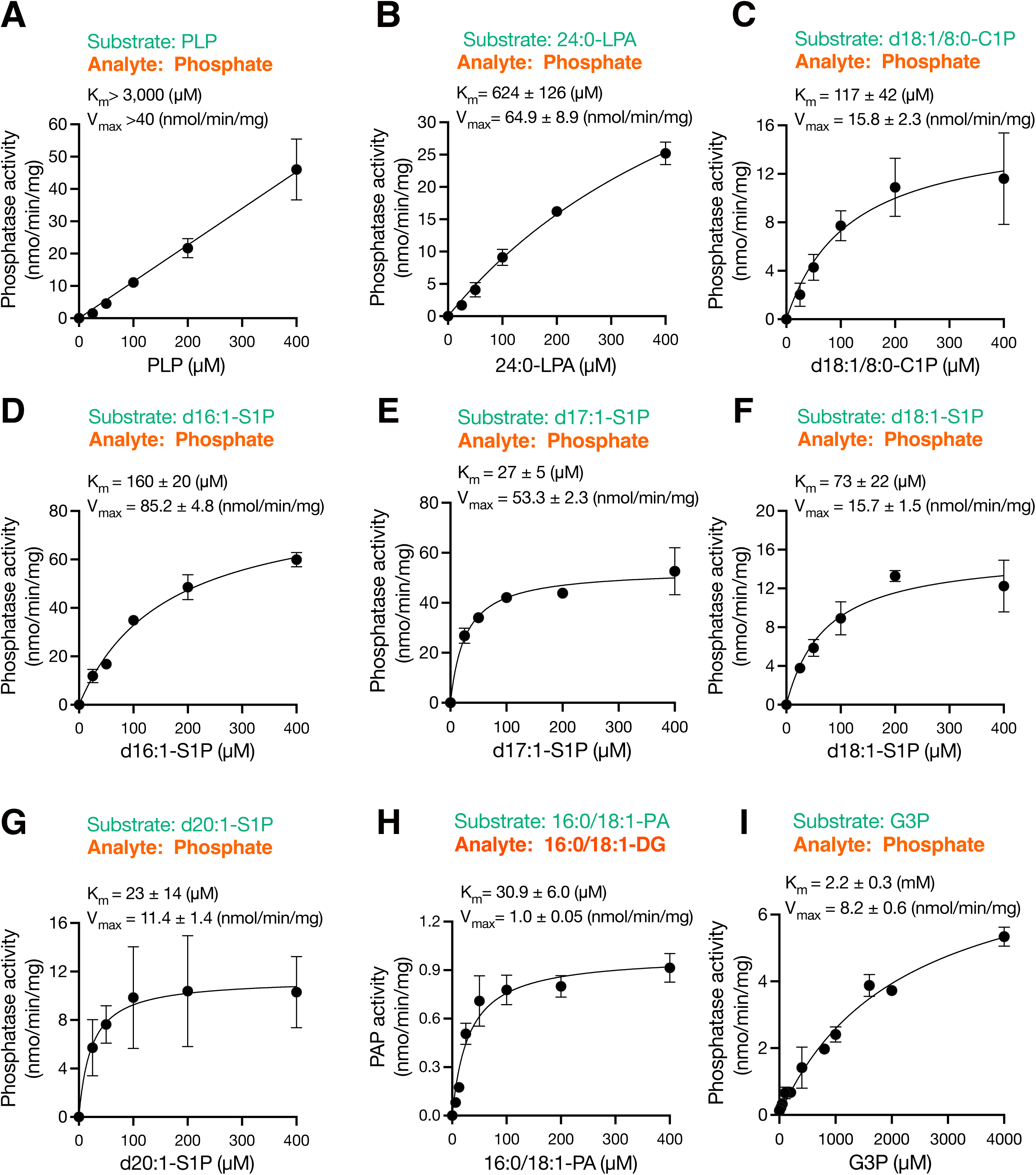
Kinetic analysis of phosphatase activities of PHOSPHO2. Phosphatase activities of PHOSPHO2-TS were measured using BIOMOL Green reagent or LC-MS/MS in the presence of various concentrations of substrates: PLP (A), 20:4-LPA (B), d18:1/8:0-C1P (C), d16:1-S1P (D), d17:1-S1P (E), d18:1-S1P (F), d20:1-S1P (G), 16:0/18:1-PA (H) and G3P (I). Values are presented as means ± S.D. (*n* = 3, technical replicates). The enzyme kinetic parameters (*K*_m_ and *V*_max_) were obtained by fitting kinetic models to the experimental data using GraphPad Prism 9.

### Effects of point mutations in PHOSPHO2 on its phosphatase activities

The amino acid sequences of PHOSPHO1 and PHOSPHO2 reportedly contain the HAD motifs that are conserved within the HAD superfamily of Mg^2+^-dependent hydrolases [28] (Fig. 8A and Suppl. Fig. S2). The catalytic core residues are highly conserved throughout the HAD phosphatase family and are clustered into four motifs: Motif I: hhhDxDx(T/V)(L/V)h (where h represents a hydrophobic residue, and x indicates any amino acid); Motif II: hhhhhh(S/T); Motif III: Lys residue, which is spaced 18–30 residues apart from motif IV; Motif IV, (G/S)(D/S)x_3-4_(D/E)hhh. Both PHOSPHO1 and PHOSPHO2 contain HAD motifs (Fig. 8A and Suppl. Fig. S2) (LVFDFDNTII, residues 5–14 of PHOSPHO2; SD, residues 98–99; K-X_24_-GDx_3_D, residues 153–183), which led us to evaluate the effects of point mutation (D8A, D99A, and D179A of PHOSPHO2) (Fig. 8B) on phosphatase activities of PHOSPHO2 (Fig. 8C–F). Substitution of the Asp8 and Asp179 with Ala (D8A or D179A) abolished PLP-, LPA-, and S1P-phosphatase activity (> 99% decrease) and strongly decreased PAP activity (over 80% decrease). PHOSPHO2^D99A^ exhibited weak phosphatase activities toward PLP, LPA, and S1P (58–75% decreased) compared to wild-type PHOSPHO2. However, PAP activity was not changed by the mutation (D99A). These results suggest that Asp8 and Asp179 residues are crucial for the phosphatase activities of PHOSPHO2, and Asp99 weakly contributes to its activity.

**Figure 8.**
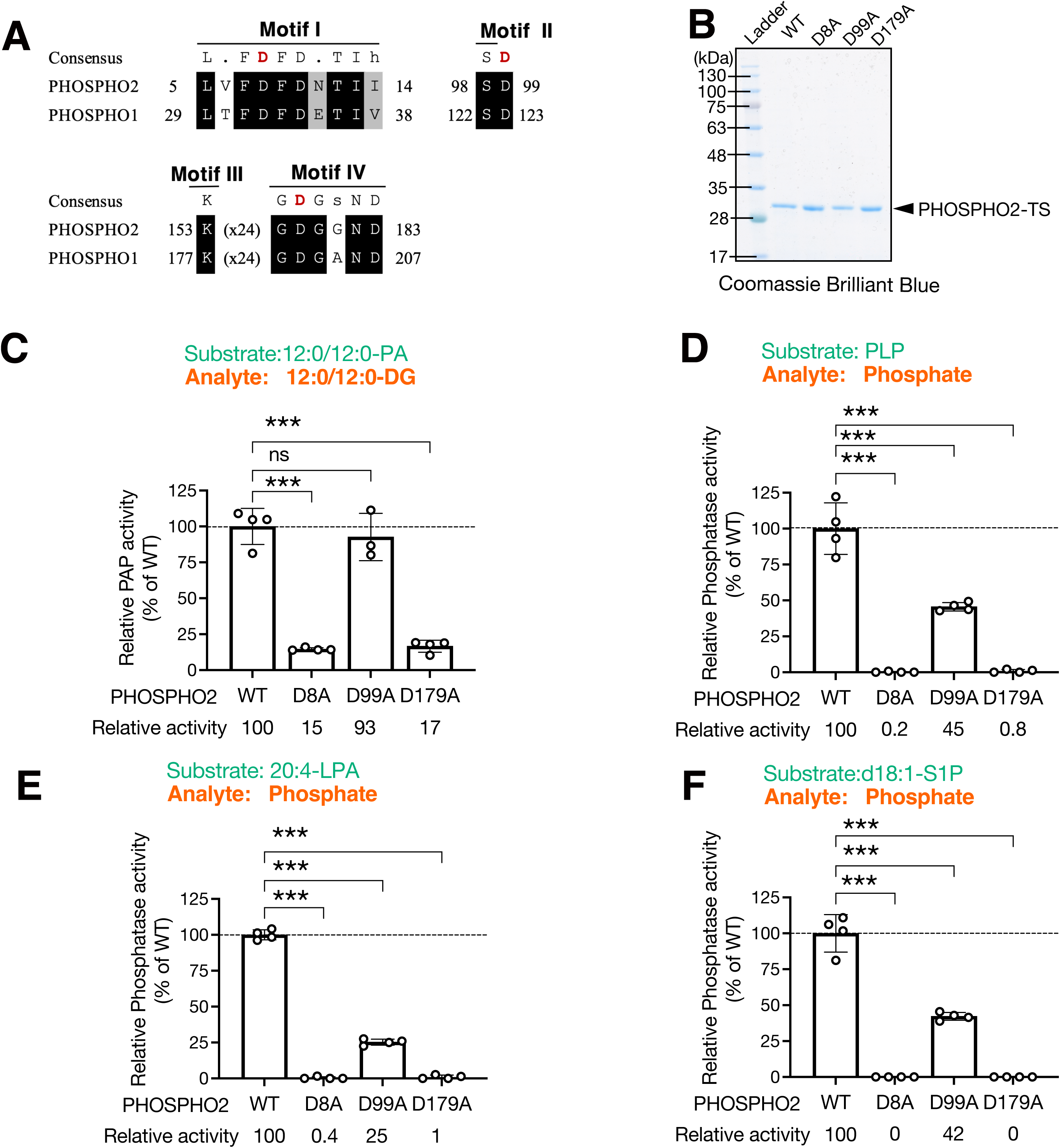
Effects of point mutations in the conserved catalytic site of PHOSPHO2 on phosphatase activities. (A) Sequence alignment of the active site regions of human PHOSPHO1 (NCBI Reference Sequence: NP_848595.1, UniProt accession number: Q8TCT1) and PHOSPHO2 (NCBI Reference Sequence: NP_001008489.1, UniProt accession number: Q8TCD6) was created using ClustalW (version 2.1) provided from the DNA Data Bank of Japan (DDBJ) [88]. (*B*) Wild-type human PHOSPHO2-TS (WT) and PHOSPHO2 point mutants in conserved catalytic motifs (D8A, D99A, and D179A) [28] were expressed in *E. coli* BL21 cells and purified. Purified proteins were detected using SDS-PAGE followed by Coomassie Brilliant Blue staining. BlueStar Prestained Protein Ladder was used as a molecular weight marker. (*C*–*F*) Effects of the point mutations on phosphatase activities toward PA (*C*), PLP (*D*), LPA (E), and S1P (*F*) were examined. Relative phosphatase activities were shown compared to activity of wild-type PHOSPHO2 (set at 100%). Data are presented as means ± S.D. (*n* = 4, technical replicates). ***, *p* < 0.005. ns, not significant. One-way ANOVA with Dunnett’s post hoc test was used.

To examine the effect of point mutant (D8A, D99A, and D179A) on the pocket formation of PHOSPHO2, the predicted structures of PHOSPHO2 point mutants were generated using AlphaFold3 [46] and their putative substrate-binding pocket volumes were calculated using the Computed Atlas of Surface Topography of the universe of protein Folds (CASTpFold) server [47] (Table3). The putative pocket volume of PHOSPHO2^D99A^ was larger than that of the wild type and other point mutants. These results suggest that the pocket structure of PHOSPHO2^D99A^ was altered, resulting in lower phosphatase activity compared to the wild-type PHOSPHO2.

### Effects of NEM and propranolol on phosphatase activities of PHOSPHO2

We determined the effects of the inhibitors on other phosphatase activities (Table 2 and Fig. 9). In contrast to PAP activity (Fig. 5A), phosphatase activities toward PLP, LPA, and S1P were not affected by propranolol. C1P phosphatase activity was inhibited in the presence of higher concentrations of propranolol (over 500 µM). These results suggested that propranolol is a PAP selective inhibitor for PHOSPHO2.

**Figure 9.**
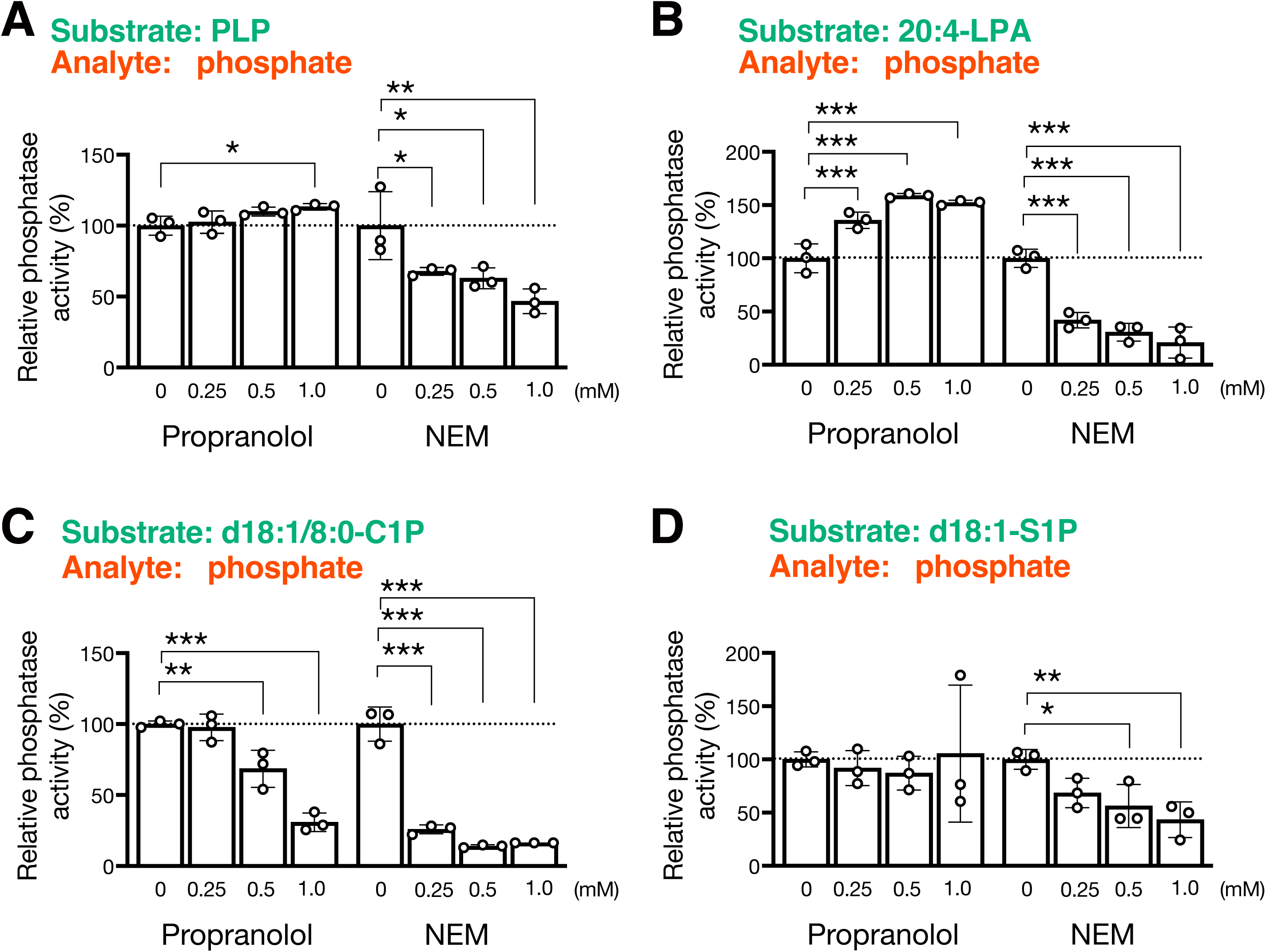
Effects of NEM and propranolol on phosphatase activities of PHOSPHO2. In the presence of various concentrations of *N*-ethylmaleimide (NEM) or propranolol, phosphatase activities of PHOSPHO2 toward PLP (A), LPA (B), C1P (C), and S1P (D) were measured. Values are presented as percentages of the activity of PHOSPHO2 in the absence of inhibitor (set at 100%). Values are presented as means ± S.D. (*n* = 3, technical replicates). *, *p* < 0.05; **, *p* < 0.01; ***, *p* < 0.005 (*vs.* control (without inhibitor)). One-way ANOVA with Dunnett’s post hoc test was used.

**Table 2.** Kinetic constants for *K*m and *V*max of PHOSPHO2 and effects of inhibitors and lipids on the phosphatase activity.

| Substrate | Molecular species | $K_m$<br>($\mu M$) | $V_{max}$<br>(nmol/min/mg) | NEM | Propranolol | Phospholipid<br>(PC and PE) | Other inhibitor<br>(lipid) |
| --- | --- | --- | --- | --- | --- | --- | --- |
| PLP |  | > 3,000 | > 40 | ++ | – | – | PI |
| PA | 16:0/18:1 | 30.9 ± 6.0 | 1.0 ± 0.05 | ++ | ++ | Increased | PI, Sph, LPA, Ceramide |
| | 12:0/12:0 | 798 ± 102 | 10.5 ± 1.0 | | | | |
| LPA | 20:4 | 624 ± 126 | 64.9 ± 8.9 | ++ | – | – | PI, Sph |
| C1P | d18:1/8:0 | 117 ± 42 | 15.8 ± 2.3 | ++ | + | – | Sph |
| S1P | d16:1 | 160 ± 20 | 85.2 ± 4.8 | + | – | – | |
| | d17:1 | 27 ± 5 | 53.3 ± 2.3 | | | | |
| | d18:1 | 73 ± 22 | 15.7 ± 1.5 | | | | |
| | d20:1 | 23 ± 14 | 11.4 ± 1.4 | | | | |
| G3P | | 2,200 ± 300 | 8.2 ± 0.6 | NT | NT | Inhibited | |

The $K_m$ and $V_{max}$ were calculated from the data (Figs. 4 and 7).
+, phosphatase activity was inhibited in the presence of the inhibitor (1.0 mM)
++, phosphatase activity was inhibited in the presence of the inhibitor (0.25 mM)
–, not changed
NT, not tested

Conversely, NEM inhibited all phosphatase activities of PHOSPHO2 including PAP activity, indicating that NEM is pan-phosphatase inhibitor of PHOSPHO2.

### Effects of various lipids on the phosphatase activities of PHOSPHO2

We determined whether lipids affect the phosphatase activities of PHOSPHO2 (Table 2 and Fig. 10). C1P and LPA phosphatase activities were significantly inhibited by Sph. PI inhibited PLP and LPA phosphatase activities. Other lipids (PC, PE, DG, MG, Cer, and Sphinganine) were not observed to have any effect on phosphatase activities. Notably, PC and PE, which increased the PAP activity (Fig. 5B), did not increase other phosphatase activities of PHOSPHO2, indicating that PHOSPHO2 preferentially exhibits PAP activity on cellular membranes where PC and PE are enriched.

**Figure 10.**
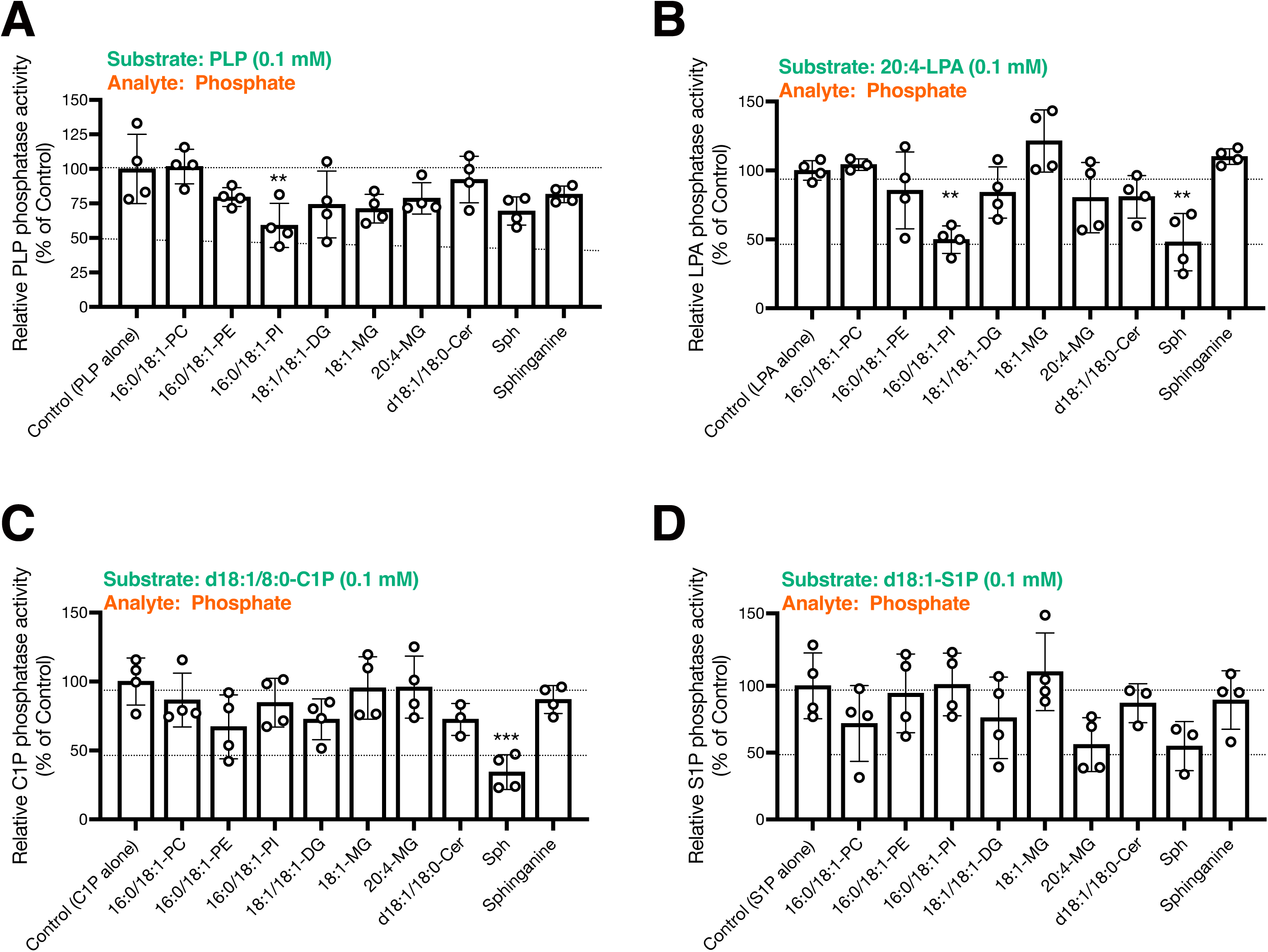
Effects of various lipids on phospholipid phosphate phosphatase activities of PHOSPHO2. Phosphatase activities of PHOSPHO2 toward 0.1 mM of PLP (A), LPA (B), C1P (C), and S1P (D) were measured in the presence of various lipids (each at 0.1 mM). Values are presented as percentages of activity of PHOSPHO2 in the presence of the substrate alone (set at 100%). Values are presented as means ± S.D. (*n* = 3, technical replicates). *, *p* < 0.05; **, *p* < 0.01; (*vs.* control (without inhibitor)). One-way ANOVA with Dunnett’s post hoc test was used.

### Structural analysis of PHOSPHO2 using AlphaFold3

To interpret the experimental data obtained in this study, structural analyses were carried out using AlphaFold3 and the docking simulation tool including SwissDock [48] and Webina [49] (Figs. 11–13, and Suppl. Figs. S6–7). Because experimentally resolved structures of human PHOSPHO2 is currently unavailable, AlphaFold3-predicted models containing a single magnesium ion were employed (Fig. 11A). The Rossmann-like fold that is composed of at least five parallel strands in a 54123 strand order, sandwiched by α-helices at both sides, is conserved in the HAD phosphatases [11]. This typical feature was also observed in the predicted structure of PHOSPHO2 (Fig. 11B). The Mg^2+^ ion formed coordinate bonds with three aspartate residues in HAD-like motifs (Figs. 11C): side chain of N-terminal Asp of Motif-I, main-chain carbonyl of C-terminal Asp of Motif-I, and side chain of Asp of Motif -IV (D8, D10, and D179 of PHOSPHO2, respectively). The four catalytic HAD motifs conserved in PHOSPHO2 are located near the Rossman-like fold: Motif I (8-DFDNT-12) at the end of strand I; motif II (98-SD-99) at the end of strand II; motif III (K153) at the N-terminus of the helix preceding strand IV (α3 in Fig. 11B); motif IV (179-DGGND-183) between the end of strand IV and N terminus of the helix (α4). These features of the predicted PHOSPHO2 structure suggest that the AlphaFold3-predicted structure is likely accurate and can be used to interpret experimental data.

**Figure 11.**
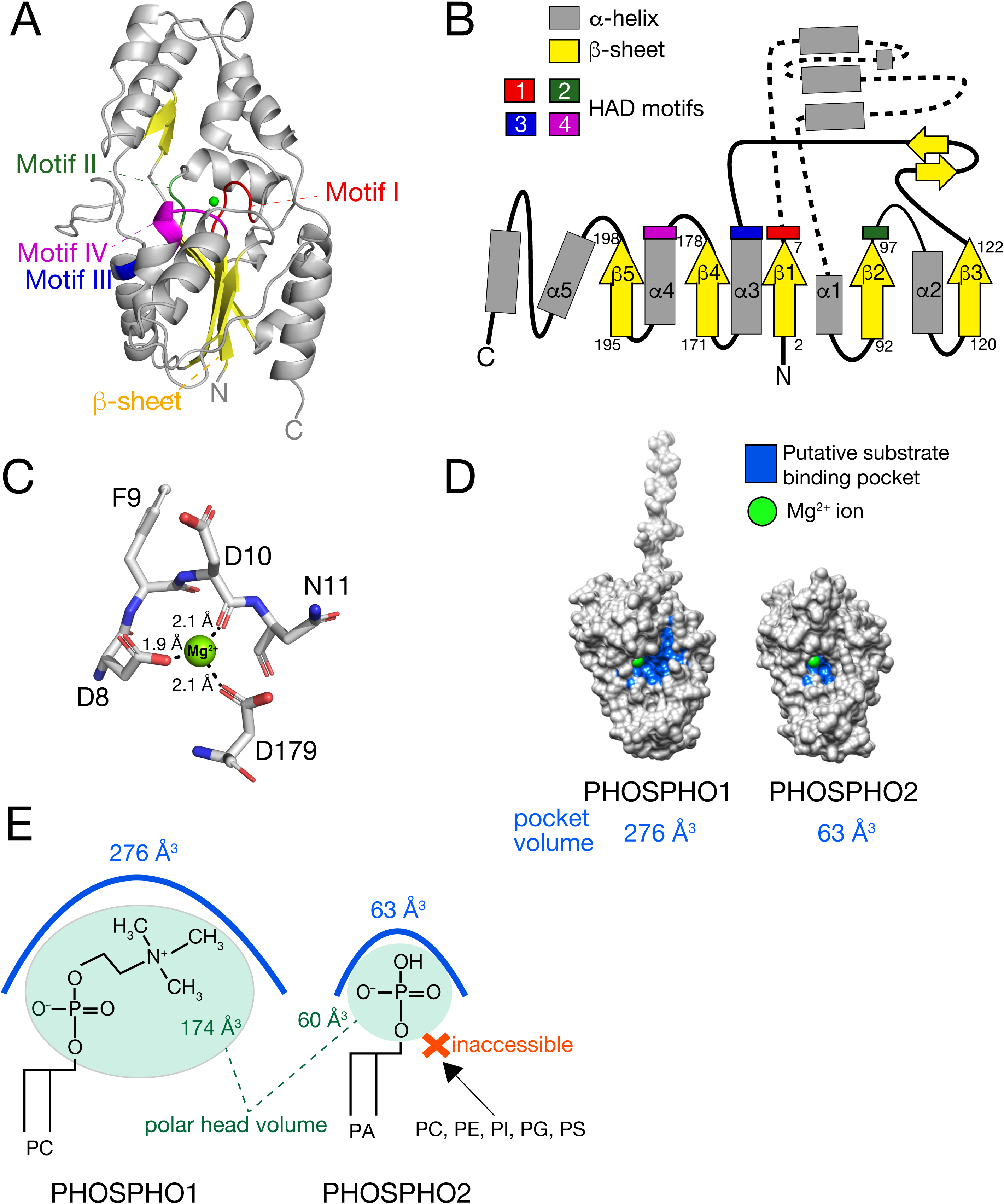
Structure analysis of AlphaFold3-predicted PHOSPHO1 and PHOSPHO2. (A) The structure of phosphatase orphan 2 (Gene name: PHOSPHO2, NCBI Gene ID: 493911, UniProt accession number: Q8TCD6) with Mg^2+^ ion was predicted using AlphaFold3 [46]. β-strands (yellow), Motif I (DFDFT, residues 8–12) (red), Motif II (S98 and D99) (green), Motif III (K153) (blue), and Motif IV (DGGND, residues 179–183) (magenta) are highlighted. Mg^2+^ is represented by a light green sphere. (B) Topology diagram of PHOSPHO2. β-strands and α-helices are numbered sequentially in the order of their appearance in the primary sequence. All colors correspond to the panel (A). (C) Contacts between Mg^2+^ ion and PHOSPHO2 were analyzed using UCSF ChimeraX (version 1.10) [89] and visualized using PyMOL. Dotted lines indicate coordination bonds. Carbon (grey), nitrogen (blue), and oxygen (red) are highlighted. The Mg^2+^ ions are represented as a green sphere. (D) Comparison of the substrate binding pocket of PHOSPHO1 and PHOSPHO2. Surface pockets in the AlphaFold3-predicted structures were predicted and visualized using CASTpFold [47] and UCSF Chimera (version 1.19) [89]. Putative pockets are shown in blue. The Mg^2+^ ion is represented by a light green sphere. Note that the catalytic Asp residues in Motif I and Motif IV (D8, D10, and D179 of PHOSPHO2) contribute to pocket formation. (E) Schematic model of substrate selectivity of PHOSPHO1 and PHOSPHO2. Pocket volumes were calculated using CASTpFold (Table 3). Molecular volumes (Van der Waals volume + Probe excluded void volume) were calculated using MoloVol [90]. Calculated volumes (Å^3^) are as follows: PO_4_, 60.4; PCHO, 173.6; PEA, 121.2; Phospho-L-serine, 151.2; Inositol phosphate, 216.7; *sn*-Glycerol 3-phosphate, 144.2; H_2_O, 19.3.

**Figure 12.**
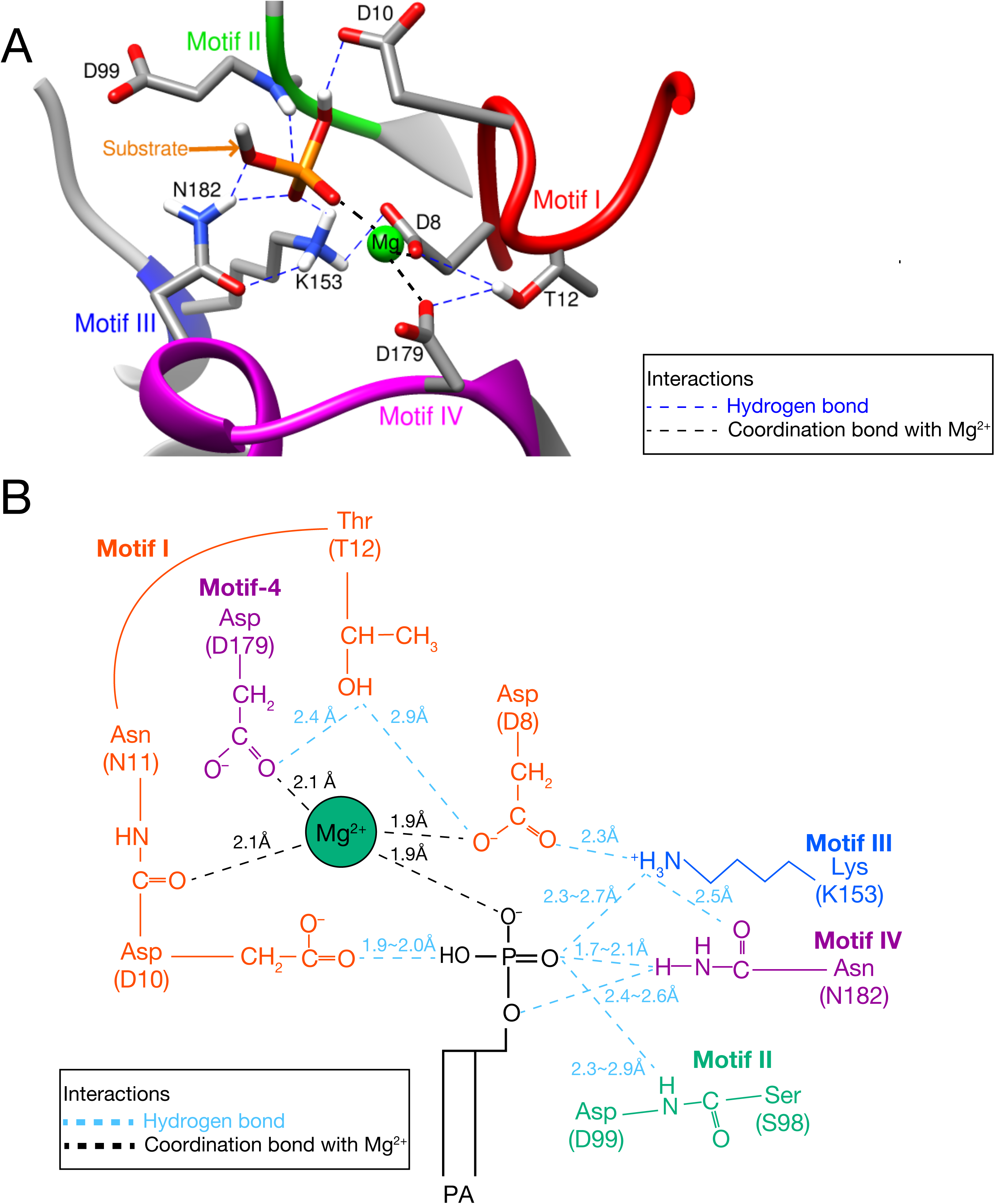
Active site of AlphaFold3-predicted PHOSPHO2 docked with PA. (A) Detailed view of active site of PHOSPHO2 with Mg^2+^ and docked substrate. In this model, 3:0/3:0-PA was used for the docked substrate. The docking model was generated using SwissDock [48]. Models were visualized and analyzed using UCSF Chimera [89]. The representative docked structure with the lowest binding free energy (ΔG (kcal/mol)) is shown. For clarity, the glycerol backbone of PA is omitted from the figure. Colored spheres indicate atoms: carbon (orange), nitrogen (blue), oxygen (red), hydrogen (white), and Mg^2+^ (light green). Colored dotted lines indicate hydrogen bonds (blue) or Mg^2+^ coordination bonds (black). (B) A schematic representation of active site of PHOSPHO2 docked with PA. Three independent docking runs were performed, and the binding mode with the lowest binding free energy (ΔG) was selected for analysis. Bond distances (hydrogen bonds and Mg^2+^-coordination bonds) were measured using PyMOL.

**Figure 13.**
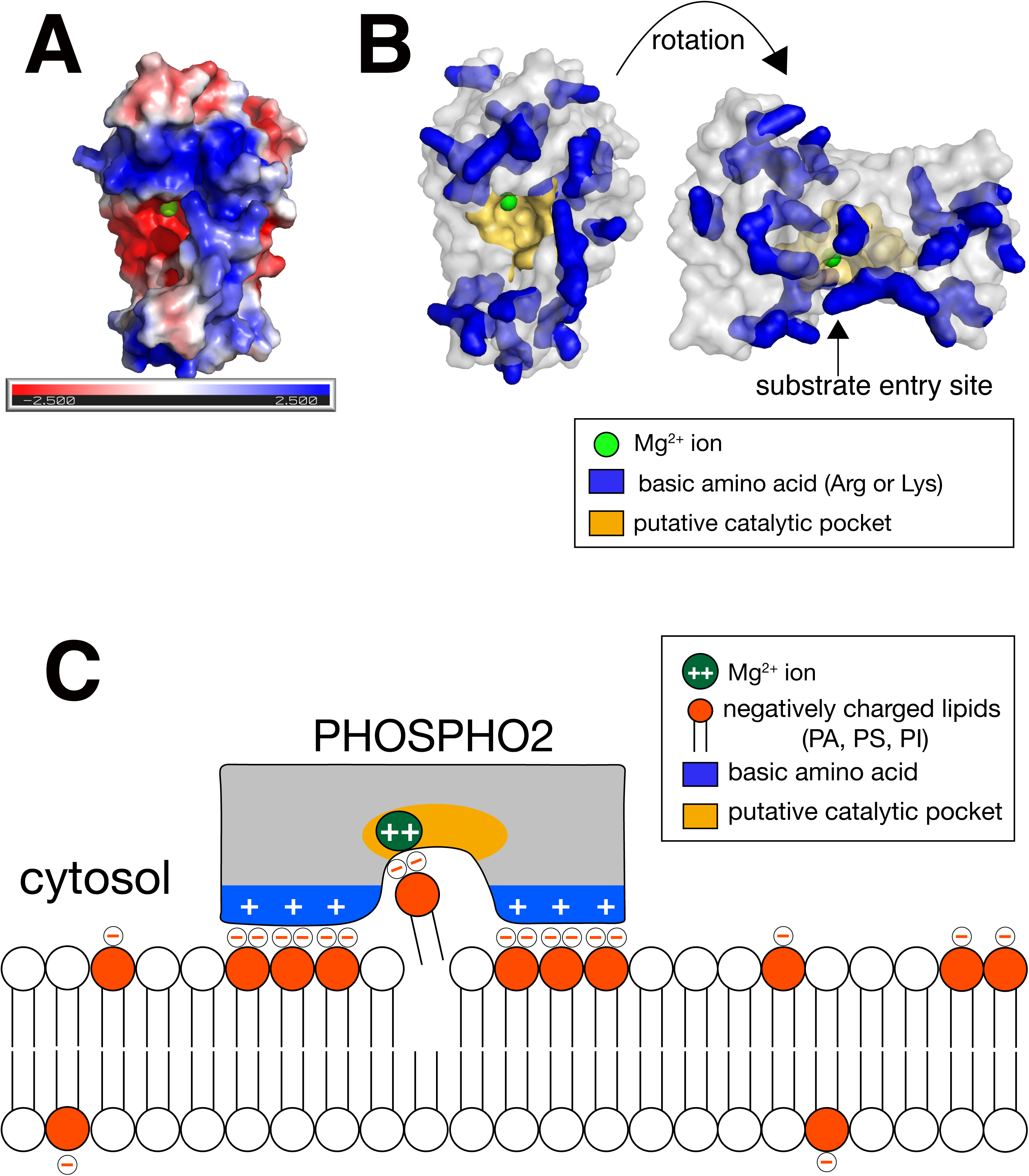
Electrostatic potentials of AlphaFold3-predicted PHOSPHO2. (A) The electrostatic potential was calculated using PyMOL and APBS (Adaptive Poisson-Boltzmann Solver). Positive and negative surfaces are shown in blue and red, respectively. The Mg^2+^ ion is represented as a light green sphere. (B) Positively charged amino acids (Lys and Arg residues) of PHOSPHO2 are visualized. These basic residues are shown in blue. Putative substrate-binding pockets predicted by CASTpFold [47] (Fig. 11D and Table 3) are shown in yellow. The Mg^2+^ ion is represented as a light green sphere. (C) A model for the interaction of PHOSPHO2 with PA on the plasma membrane. The cytosolic leaflets of membranes including the plasma, Golgi and endosomal membranes are negatively charged due to the enrichment of anionic phospholipids such as PS and PI/phosphoinositides (the net charge: –1) [91]. Upon interaction with Lys or Arg residues, the net charge of PA increases from –1 to –2 and this change stabilizes protein-lipid interactions (the electrostatic/hydrogen bond switch model) [83]. Because Arg and Lys residues are enriched around the substrate entry site of PHOSPHO2, the protein-lipid interaction may be stabilized by increased charge of PA by the positively charged surface amino acids of PHOSPHO2.

**Table 3.** Quantification of substrate binding pockets calculated with CASTpFold.

| Enzyme | SA area<sup>1</sup> | SA volume<sup>2</sup> | Mouth SA area<sup>3</sup> | SA circumference sum<sup>4</sup> | Amino acid residue Contributing pocket<sup>5</sup> |
| --- | --- | --- | --- | --- | --- |
| PHOSPHO1 | 288.2 | 276.0 | 86.0 | 69.1 | **Asp32**, **Asp34**, Glu35, Asn41, Asp43, Tyr63, Glu65, Gly66, Tyr68, Ser122, **Asp123**, Ala124, Ala169, Arg170, Cys171, Pro172, **Lys177**, **Asp203**, Gly204, Ala205, Asn206, Asp207, Phe208, Cys209 |
| PHOSPHO2 Wild type | 118.5 | 62.6 | 28.4 | 31.6 | **Asp8**, Phe9, **Asp10**, Asn11, Asn17, Ser18, Asp19, Trp44, Ser98, **Asp99**, Ser100, **Lys153**, **Asp179**, Gly180, Asn182, Asp183 |
| PHOSPHO2 D8A | 97.2 | 46.9 | 23.7 | 29.0 | **Ala8**, Phe9, **Asp10**, Asn17, Ser18, Asp19, Trp44, Ile97, Ser98, **Asp99**, Ser100, **Lys153**, **Asp179**, Asn182, Asp183 |
| PHOSPHO2 D99A | 658.1 | 615.2 | 169.4 | 95.9 | **Asp8**, Phe9, **Asp10**, Asn11, Asn17, Ser18, Asp19, Thr20, Lys41, Gly42, Phe43, Trp44, Thr45, Glu48, Phe52, Ser98, **Ala99**, Ser100, Asn101, Phe104, Asn124, Val136, Asn138, Arg146, Cys147, Pro148, Lys149, Asn150, Leu151, Cys152, **Lys153**, Lys154, **Asp179**, Gly180, Gly181, Asn182, Asp183, Val184, Cys185, Pro186, Tyr202, Thr203, Leu204, Thr207, Arg210 |
| PHOSPHO2 D179A | 120.3 | 62.2 | 28.1 | 31.1 | **Asp8**, Phe9, **Asp10**, Asn11, Thr12, Asn17, Ser18, Asp19, Trp44, Ser98, **Asp99**, Ser100, **Lys153**, **Ala179**, Gly180, Asn182, Asp183 |

<sup>1</sup> pocket solvent-accessible surface area (Å<sup>2</sup>).
<sup>2</sup> pocket volume based on the solvent-accessible surface (Å<sup>3</sup>).
<sup>3</sup> total area of mouth opening based on the solvent-accessible surface (Å<sup>2</sup>).
<sup>4</sup> total circumference of mouth opening based on the solvent-accessible surface (Å).
<sup>5</sup> residues colored in red are part of HAD motifs (Suppl. Fig. S2).

### Comparison of the putative substrate-binding pockets of PHOSPHO1 and PHOSPHO2

Using the AlphaFold3-predicted PHOSPHO1 and PHOSPHO2 structures, the substrate-binding pockets were predicted using CASTpFold server [47] (Fig. 11D and Table 3). The prediction tool detected a cavity near the catalytic Asp (Motif I) and Mg^2+^ ion in both PHOSPHO2 and PHOSPHO1. The substrate-binding pocket volume of PHOSPHO1 (276 Å^3^) is larger than the volumes of phosphocholine (173.6 Å^3^), and phosphoethanolamine (121.2 Å^3^), whereas that of PHOSPHO2 (62.6 Å^3^) is comparable to that of PO_4_ (60.4 Å^3^), suggesting that PHOSPHO2 is unable to hydrolyze phospholipids with large polar head groups (PC, PE, PS, PG, and PI) due to its small substrate-binding pocket (Fig. 11E).

### Insights into the phosphatase reaction of PHOSPHO2 using AlphaFold3 and the molecular docking simulations

To predict the possible binding position of the AlphaFold3-predicted PHOSPHO2 structure and phosphate compounds, molecular docking simulations were carried out using SwissDock [48] (Fig. 12). The pose with the lowest binding free energy (ΔG) (strongest binding) was selected for the analysis. All phosphate compounds were positioned within the putative catalytic pocket. Moreover, the docked substrate formed hydrogen bonds with the HAD motif of PHOSPHO2, and formed coordination bonds with Mg^2+^ (Fig. 12B). Furthermore, substitution of Asp8 with Ala in PHOSPHO2 decreased the PLP-, LPA-, S1P-, and PA phosphatase activities (Fig. 8). These results suggest that phosphorus compounds are hydrolyzed by catalytic Asp residues (D8 and D10) via the same reaction mechanism as the hydrolase of the HAD phosphatase (Suppl. Fig. S1A) [9–11]. The D179 of PHOSPHO2 is not a catalytic Asp residue (Suppl. Fig. S1A), but it was also essential for the enzyme activity (Fig. 8). The docking model shows that D179 forms a coordination bond with the Mg^2+^ ion (Fig. 12B). This coordination bond may be essential for the enzyme activity.

### Effects of acyl chains of phospholipids on substrate–enzyme binding free energies

The predicted substrate–enzyme binding free energies (ΔG) are shown in Table 4. Compared to PLP, other substrates such as PA, LPA, S1P, and C1P can form a complex with PHOSPHO2 with ΔG values comparable to that of PLP. These data suggest that PHOSPHO2 can form a stable complex with phosphate-containing lipids and thus PHOSPHO2 may hydrolyze phospholipids in addition to PLP. To evaluate the effect of acyl chain of phosphate-containing lipids on the binding affinity with PHOSPHO2, several LPA species were docked with PHOSPHO2 (Table 4). PUFA-containing LPAs (20:4, 18:3, and 18:2) exhibited the lowest ΔG values. However, trans-unsaturated fatty acid-containing LPA showed higher ΔG values comparable to those of saturated fatty acid, indicating that only cis double bonds in fatty acids contribute to the stability of binding with PHOSPHO2. Among SFA-containing LPA molecular species, 14:0-LPA showed the lowest ΔG. This tendency in ΔG values correlates with the experimental enzyme activities (Fig. 6B).

**Table 4.**
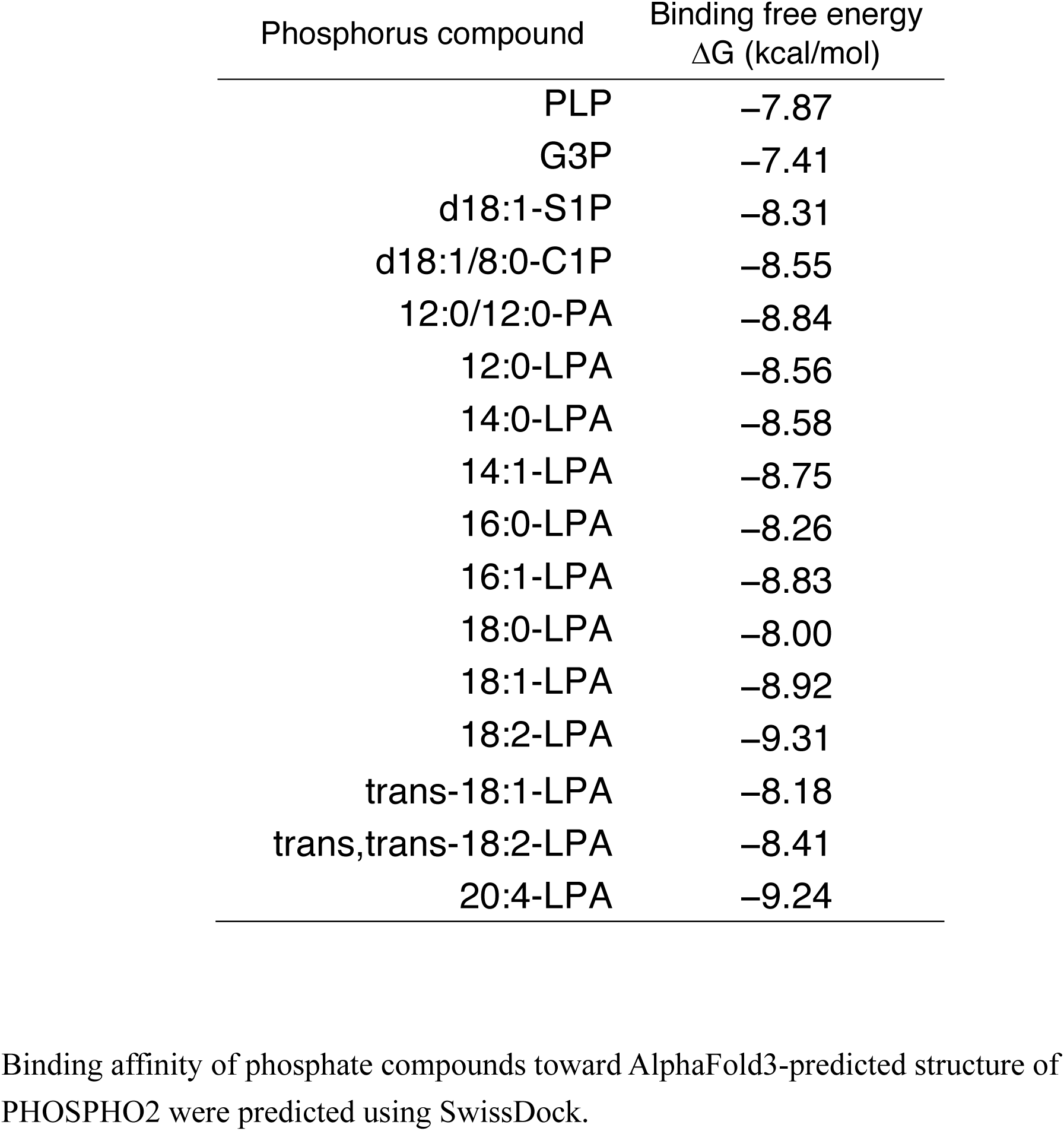
Estimated binding free energy of the docked PHOSPHO2-phosphorus compound complex.

| Phosphorus compound | Binding free energy<br>$\Delta G$ (kcal/mol) |
| --- | --- |
| PLP | −7.87 |
| G3P | −7.41 |
| d18:1-S1P | −8.31 |
| d18:1/8:0-C1P | −8.55 |
| 12:0/12:0-PA | −8.84 |
| 12:0-LPA | −8.56 |
| 14:0-LPA | −8.58 |
| 14:1-LPA | −8.75 |
| 16:0-LPA | −8.26 |
| 16:1-LPA | −8.83 |
| 18:0-LPA | −8.00 |
| 18:1-LPA | −8.92 |
| 18:2-LPA | −9.31 |
| trans-18:1-LPA | −8.18 |
| trans,trans-18:2-LPA | −8.41 |
| 20:4-LPA | −9.24 |

Binding affinity of phosphate compounds toward AlphaFold3-predicted structure of PHOSPHO2 were predicted using SwissDock.

### Analysis of protein surface properties of PHOSPHO2

We calculated electrostatic potential of the AlphaFold-generated PHOSPHO2 structure using the Adaptive Poisson-Boltzmann Solver (APBS) in Pymol (Fig. 13A). Region around the substrate entry site of PHOSPHO2 is positively charged because the Lys and Arg residues are abundant around the substrate entry site (Fig. 13B). Because polar head of PA is just a phosphomonoester, PA is negatively charged in cell. Therefore, the electrostatic potential of predicted PHOSPHO2 appears well suited for PA binding (Fig. 13C).

## Discussion

### PHOSPHO2 is defined as a new type of Mg^2+^-dependent PLPP

Because PA, LPA, S1P, and C1P are bioactive lipids, PLPPs play important roles in signal transduction of a wide variety of physiological events and diseases such as glycerolipid biosynthesis, vascular development, tumorigenesis, coronary disease, and adipogenesis [8]. PLPP genes (PAP2/LPP) have been cloned as genes encoding Mg^2+^-independent, NEM-insensitive, and transmembrane PLPPs. LPIN genes encode cytosolic Mg^2+^-dependent PAP, which hydrolyze only PA [17]. NEM-sensitive C1P phosphatase and LPA phosphatase activities were detected in the cytosolic fraction [22, 23]. However, NEM-sensitive cytosolic PLPPs have not been identified to date.

In the present study, we purified PHOSPHO2 from soluble fractions with high purity (>90% purity) (Fig. 1 and Suppl. Fig. S3A). We demonstrated for the first time that purified PHOSPHO2 exhibits Mg^2+^-dependent (Fig. 3D), NEM-and propranolol-sensitive PAP activity *in vitro* (Figs. 1D, 1G, and 5A). Moreover, PHOSPHO2 displayed higher PAP activity toward medium-chain fatty acid-containing PA and PUFA-containing PA species (Fig. 2). Furthermore, PHOSPHO2 also hydrolyzed LPA, C1P, S1P, and G3P in addition to PA and PLP (Fig. 6A). Taken together, PHOSPHO2 may act as a novel type of PLPP with enzymological properties similar to those of PAP1, but displays phospholipid phosphatase activity (EC 3.1.3.113) similar to that of PLPPs (PAP2/LPP). Therefore, PHOSPHO2 is a promising candidate for the long-sought NEM-sensitive cytosolic PLPP.

PHOSPHO2 has been identified by homology search for homologous proteins to PHOSPHO1, and it has been demonstrated that PHOSPHO2 displays PLP phosphatase activity. However, its PLP activity is extremely low compared to the major enzyme activity of PHOSPHO1 (approximately 10-fold lower than the PEA/PCho phosphatase activity of PHOSPHO1) [28]. Moreover, the biological significance of PHOSPHO2 in vitamin B6 metabolism has not been reported to date. Recently, we showed that PHOSPHO1 exhibits PC-phospholipase C (PC-PLC) (EC 3.1.4.3) and PE-PLC (EC 3.1.4.62) activities in addition to PEA/PCho phosphatase activity (EC 3.1.3.75) [38]. These findings led us to hypothesize that PHOSPHO2 may also hydrolyze phospholipids.

### Similarities and differences between PHOSPHO2 and other PAPs

The enzymological properties and biological significance of PHOSPHO2 remain largely unknown. Recent studies have shown that high mRNA expression of PHOSPHO2 is associated with the development of hepatocellular carcinoma [34, 35], and tumorigenic processes in gastric cancer cell lines [36], indicating that PHOSPHO2 plays important roles in carcinogenesis. Further studies are needed to elucidate the carcinogenesis-related functions of PHOSPHO2 as a phospholipid phosphatase enzyme (EC 3.1.3.113) *in vivo*.

The optimum enzyme reaction conditions for phospholipid phosphatase activity of PHOSPHO2 were as follows: around 40°C (Fig. 3A), pH 7.5 (Fig. 3C), 1 mM Mg^2+^ (Fig. 3E). The total intracellular Mg^2+^ concentration ranges from 5 to 20 mM [50]. However, intracellular free Mg^2+^ is approximately 0.25–1.0 mM [51] because most Mg^2+^ forms complexes with nucleotides (e.g., ATP, DNA) or other Mg^2+^-binding proteins. Under *in vitro* experimental conditions, PHOSPHO2 displayed PAP activity at 0.1 mM or higher concentrations of Mg^2+^ (Fig. 3E). Taken together, these findings indicate that PHOSPHO2 can exhibit PLPP activity in cells.

The reaction of PA dephosphorylation by PHOSPHO2 obeyed the surface dilution kinetic model [45] (Suppl. Fig. S4). The *K*_m_ value of the PAP activity was approximately 3 mol%. This value is comparable to that of Lipin-1 and PLPPs (Table 1). Moreover, the concentration of PA on the surface of the cellular membrane is approximately 1–2 mol% of total lipids [52, 53]. Taken together, it is likely that PHOSPHO2 acts as PAP on mammalian cellular membranes.

The biochemical characteristics of PHOSPHO2 as a PAP observed in the present study are similar to those of the mammalian Mg^2+^-dependent PAP previously reported [17, 54–57]. For instance, **(A) heat stability:** mammalian Mg^2+^-dependent PAP (Type 1 PAP) was more sensitive to heat treatment (55°C) than Mg^2+^-independent PAP (Type 2 PAP) [55]. The PAP activity of PHOSPHO2 was inactivated by incubation at 50°C (50% decrease) (Fig. 3B). **(B) optimal pH**: optimal pH for mammalian PAPs is around pH 7 (with a broad optimal range between pH 6.5 and 8.0) [17]. The optimum pH for PHOSPHO2 was similarly broad, between pH 6.0 and 8.0 (Fig. 3C). **(C) molecular weight:** cytosolic Mg^2+^-dependent PAP activity was detected in fractions with a molecular weight of approximately 40 kDa and 10 kDa [57]. Because the molecular mass of PHOSPHO2 is 27.8 kDa, PHOSPHO2 could be contained in the fractions that exhibited Mg^2+^-dependent PAP [56]. **(D) optimum Mg^2+^ concentration:** It has been reported that a free Mg^2+^ concentration of approximately 2 mM is optimal for mammalian PAP activity, whereas higher Mg^2+^ (approximately 10 mM) inhibits this activity [54, 55]. The same tendency was observed with purified PHOSPHO2 (Fig. 3E). **(E) inhibitors:** Several inhibitors of mammalian type 1 PAP have been reported: NEM, propranolol, divalent metal ions (Mn^2+^, Zn^2+^, Ca^2+,^ and Co^2+^), and Sph (Table 2) [17, 54]. The PAP activity of PHOSPHO2 was sensitive to these PAP inhibitors (Fig. 3F and 5A). What are the differences between PHOSPHO2 and the already-identified PAPs (LPP/PAP2, lipin-1, and sphingomyelin synthase-related protein (SMSr, Gene name: SAMD8) [43]) (Table 2)? Lipin-1 and PHOSPHO2 share similar properties (Table 1). For instance, they are both soluble proteins conserved within the HAD superfamily of Mg^2+^-dependent hydrolases (DXDX(T/V) motif). Both Lipin-1 and PHOSPHO2 are inhibited by NEM and propranolol (Fig. 5A). Notably, Lipin-1 is more sensitive to NEM than PHOSPHO2 (IC_50_: 0.1 mM versus 0.26 mM) (Fig. 5A). Moreover, the inhibitors of Lipin-1 such as divalent metal ions (Ca^2^, Mn^2+^, and Zn^2+^) and sphingoid bases (Sph and sphinganine) also inhibited the PAP activity of PHOSPHO2 (Figs. 3F and 5B). However, there are differences between these two enzymes. First, Lipin-1 displays a broad acyl chain specificity for PA [17], whereas PHOSPHO2 exhibits higher PAP activity toward medium-chain fatty acid- and PUFA-containing PA (Fig. 2). Moreover, effects of PC, PE, and DG (feed-forward effect) (Fig. 5B) on Lipin-1 were not observed [17]. Particularly, Lipin-1 exhibits only PAP activity. Although there are similarities between Lipin-1 and PHOSPHO2, they are distinct enzymes.

### Which phospholipids does PHOSPHO2 dephosphorylate in vivo?

Which substrate (PLP, C1P, S1P, PA, LPA, G3P) would PHOSPHO2 primarily dephosphorylate *in vivo*? Based on previous reports [53, 58–60], we estimated the cellular substrate concentration (Suppl. Table S1). Next, the estimated enzyme activities under the estimated substrate concentrations [52, 59, 61–73] were calculated using these results (Figs. 4 and 7). Because PA localizes to the cellular surface, in which PC is the most abundant [53], the effects of PC on the enzyme activities of PHOSPHO2 (Figs. 5B and 10) were taken into account. Considering cellular concentrations of the substrates and kinetic constants of PHOSPHO2 (Table 2), the highest estimated enzyme activity of PHOSPHO2 in cells was PAP activity, which was at least 10-fold higher than the estimated PLP phosphatase activity (Suppl. Table 1). These results strongly suggest that, in addition to Lipin enzymes, on the cellular surface, PHOSPHO2 acts exclusively as a PAP. However, we cannot rule out the possibility that endogenous effectors may modulate the substrate selectivity of PHOSPHO2. To address the biological significances of PHOSPHO2 as a lipid-metabolizing enzyme *in vivo*, further studies are required.

PHOSPHO2 exhibited higher PAP activity toward shorter SFA- or PUFA-containing PA molecular species such as 12:0/12:0-, 14:0/14:0-, 18:0/20:4-, and 18:0/22:6-PA (Fig. 2). The acyl chain selectivity of PHOSPHO2 differs from that of other PAPs. For example, PAP2A (gene name: PLPP1) and PAP2B (gene name: PLPP3) are least active against intermediate-length SFA (14:0 and 10:0)-containing PA and are the most active against medium-chain SFA (6:0)-containing PA [44]. Lipin-1 exhibits comparable PAP activities toward 1,2-diunsaturated and 1-saturated-2-unsaturated fatty acid-containing PA. However, Lipin-1 exhibits little PAP activity toward 16:0/16:0-PA and 18:0/18:0-PA [17]. Although PHOSPHO2, PLPPs, and Lipin-1 catalyze the same reaction, they likely generate distinct PA molecular species due to differences in their acyl chain specificities, and hence these PAPs may regulate distinct signal transductions.

The tendency of the predicted substrate–enzyme binding free energies (ΔG) (Table 4) indicates that cis double bonds in fatty acids contribute to the stability of binding with PHOSPHO2. Among SFA-containing LPA molecular species, 14:0-LPA showed the lowest ΔG. These tendencies in ΔG values correlate with the experimental enzyme activities (Fig. 6B). However, there are several factors that govern enzyme activity. For example, PC and PE enhanced only the PAP activity (Fig.5B and Table 2). Notably, DG enhanced only the PAP activity (feed-forward control). Therefore, comparing predicted ΔG values alone cannot fully account for all aspects of the enzyme reactions of PHOSPHO2. However, the binding affinity of a substrate for PHOSPHO2 may, in part, contribute to the enzyme activity of PHOSPHO2 because the simulation data are not inconsistent with experimental data.

### What is the biological role of PHOSPHO2 as a PA phosphatase?

Mammalian cells contain more than 50 structurally distinct DG and PA species [3]. Recently, proteins that bind to a particular PA molecular species have been reported [4, 74]. For instance, PUFA-containing PA strongly interact with L-lactate dehydrogenase A and attenuates its enzyme activity [75]. Synaptojanin-1, a Parkinson’s disease-related protein, also selectively interacts with PUFA-containing PA, and its enzyme activity is enhanced in the presence of PA [76]. Clathrin coat assembly protein 180 (AP180), which drives endocytosis, selectively interacts with 18:0/22:6-PA, and 18:0/22:6-PA inhibits the interaction between AP180 and clathrin [77]. 18:0/22:6-PA also selectively interacts with Praja-1 E3 ubiquitin-protein ligase, which controls serotonin transporter, and activates its enzyme activity [78]. Shorter SFA-containing molecular species are also involved in biological events such as fission of vesicle formation by Coat Protein I in the late stage [79] and superoxide production [80]. Because PHOSPHO2 exhibits higher PAP activity toward PUFA- or shorter SFA-containing PA (Fig. 2), PHOSPHO2 may regulate these enzymes.

### Why do metal ions other than Mg^2+^ inhibit the PAP activity of PHOSPHO2?

Although Mg^2+^ is essential for the phosphatase activity of PHOSPHO2 (Fig. 3D), other metal ions (Mn^2+^, Ni^2+^, Ca^2+^, Co^2+^, and Zn^2^) inhibited it (Fig. 3F). For instance, the displacement of Mg^2+^ by Ca^2+^ results in the loss of activity in the HAD superfamily [81]. Based on X-ray crystal structure analysis, the inhibitory effect of Ca^2+^ on Mg^2+^-dependent phosphatases has been proposed to correlate with the differences in the coordination geometries of Ca^2+^ and Mg^2+^ in the active site. Comparing the structures of AlphaFold3-generated Ca^2+^-binding and Mg^2+^-binding PHOSPHO2 (Fig. 11C and Suppl. Fig. S6), a similar result is observed. The Asp8, Asp10, and Asp179 residues in PHOSPHO2 form coordination bonds with Mg^2+^, whereas Ca^2+^ additionally interacts with one of the oxygen atoms of Asp179 (Suppl. Fig. S6A). This additional interaction may distort the active site. As Mn^2+^ has five unpaired d-electrons, labile water exchange, and a larger ionic radius compared to Mg^2+^[82], Mn^2+^ may fail to form stable interactions with PHOSPHO2. Although the ionic radius of Zn^2+^ is close to that of Mg^2+^ [82] and Zn^2+^ forms similar coordination bonds with PHOSPHO2 (Suppl. Fig. S6C), Zn^2+^ disrupts the phosphatase activity (Fig. 3D). Compared to Mg^2+^, Zn^2+^ is a softer acid (intermediate acid), usually has coordination numbers smaller than Mg^2+^ (predominantly exhibiting a coordination number of 4, compared to 6 in Mg^2+^). These differences may affect the activity of PHOSPHO2.

### Why do PI and Sph inhibit the phosphatase activity of PHOSPHO2?

Several lipids, especially phosphatidylinositol (PI) and sphingosine (Sph), inhibited phosphatase activity of PHOSPHO2 (Table 2, Fig.5B and 10). To interpret the results, molecular docking simulation was performed (Suppl. Fig. S7A). Both PI and Sph were positioned near the entrance of putative catalytic pocket of PHOSPHO2. Sph is unlikely to be hydrolyzed by PHOSPHO2 because it lacks a phosphate ester bond. The phosphate group of PI is located far from active site of PHOSPHO2 and thus PI may not be hydrolyzed (Suppl. Fig. S7B). Therefore, these two molecules can retain the ability to bind to PHOSPHO2. These interactions may inhibit the phosphatase activity because they sterically hinder substrate access to the catalytic pocket.

### Effects of catalytic pocket volumes of PHOSPHO1 and PHOSPHO2 on their enzyme activities

Why does PHOSPHO1, but not PHOSPHO2, hydrolyze PC and PE [38] (Fig. 1D and G)? Using AlphaFold3-predicted structures, we compared structures of PHOSPHO2 and PHOSPHO1 and found that the catalytic pocket volume of PHOSPHO2 (62.6 Å^3^) is smaller than that of PHOSPHO1 (276 Å^3^) (Table 3 and Fig. 11D). Therefore, the polar head groups of PC (*i.e.* PCho, 173.6 Å^3^) and PE (*i.e.* PEA, 121.2 Å^3^) are too large to fit into the catalytic pocket of PHOSPHO2 as substrates (Fig. 11E). These predicted structural data support the biochemical results (Fig. 1D and 1G). Although AlphaFold 3-predicted structures may not fully reflect the native conformations of proteins, the difference in pocket volumes between PHOSPHO1 and PHOSPHO2 provides a structural basis for their distinct substrate specificities.

Although D99 of PHOSPHO2 is unlikely to form hydrogen bond with substrate or coordination bond with Mg^2+^ (Fig. 12B), PHOSPHO2^D99A^ displayed weaker phosphatase activity than that wild-type PHOSPHO2. The putative pocket volume of PHOSPHO2^D99A^ was larger than that of the wild type and other point mutants (Table 3), indicating that D99 may contribute to proper substrate pocket formation.

### How does PHOSPHO2 bind to PA-containing cellular membrane?

Using the AlphaFold-generated PHOSPHO2 structure, electrostatic potential on protein surfaces were visualized (Fig. 13A) The positively charged amino acids such as Lys and Arg increase the charge of PA ([HPO_3_]^−^→ PO_3_^2–^) by forming hydrogen bonds with the phosphate of PA, thereby stabilizing the protein-lipid interaction [83]. The Lys and Arg residues are abundant around the substrate entry site of PHOSPHO2 (Fig. 13B). Therefore, the electrostatic potential of predicted PHOSPHO2 appears well suited for PA binding, provides isolated hydrophobic environment (Fig. 13C) and thus PHOSPHO2 could be capable of hydrolysis of PA on the cell membrane. Moreover, negatively charged phospholipids (*e.g.* PA, PS, and PI) are enriched on the cytosolic side of the plasma membrane [84]. It is possible that PHOSPHO2 can bind to the cytosolic leaflet of the plasma membrane (Fig. 13C).

### PHOSPHO1 and PHOSPHO2 are new type of DG-generating enzymes

We previously reported that PHOSPHO1 displays PC-PLC (EC 3.1.4.3) and PE-PLC (EC 3.1.4.62) activities [38] (Table 1). Moreover, intracellular DG levels, especially SFA-containing DGs, were increased by overexpression of PHOSPHO1. Furthermore, we demonstrated that PHOSPHO1 interacted with DG kinase δ isozyme (DGKδ) (Gene name: DGKD, NCBI Gene ID: 8527, UniProt accession number: Q16760), which selectively phosphorylates SFA-containing DG [63, 85]. Unlike PHOSPHO1, PHOSPHO2 does not display PLC activity toward PC and PE (Fig. 1D and G) or interact with DGKδ [38]. This indicates that PHOSPHO2 is involved in a DG metabolic pathway distinct from that of PHOSPHO1 (proposed PHOSPHO1 pathway: PC/PE→DG→DGKδ→PA), although both PHOSPHO2 and PHOSPHO1 generate DG (Fig. 14).

**Figure 14.**
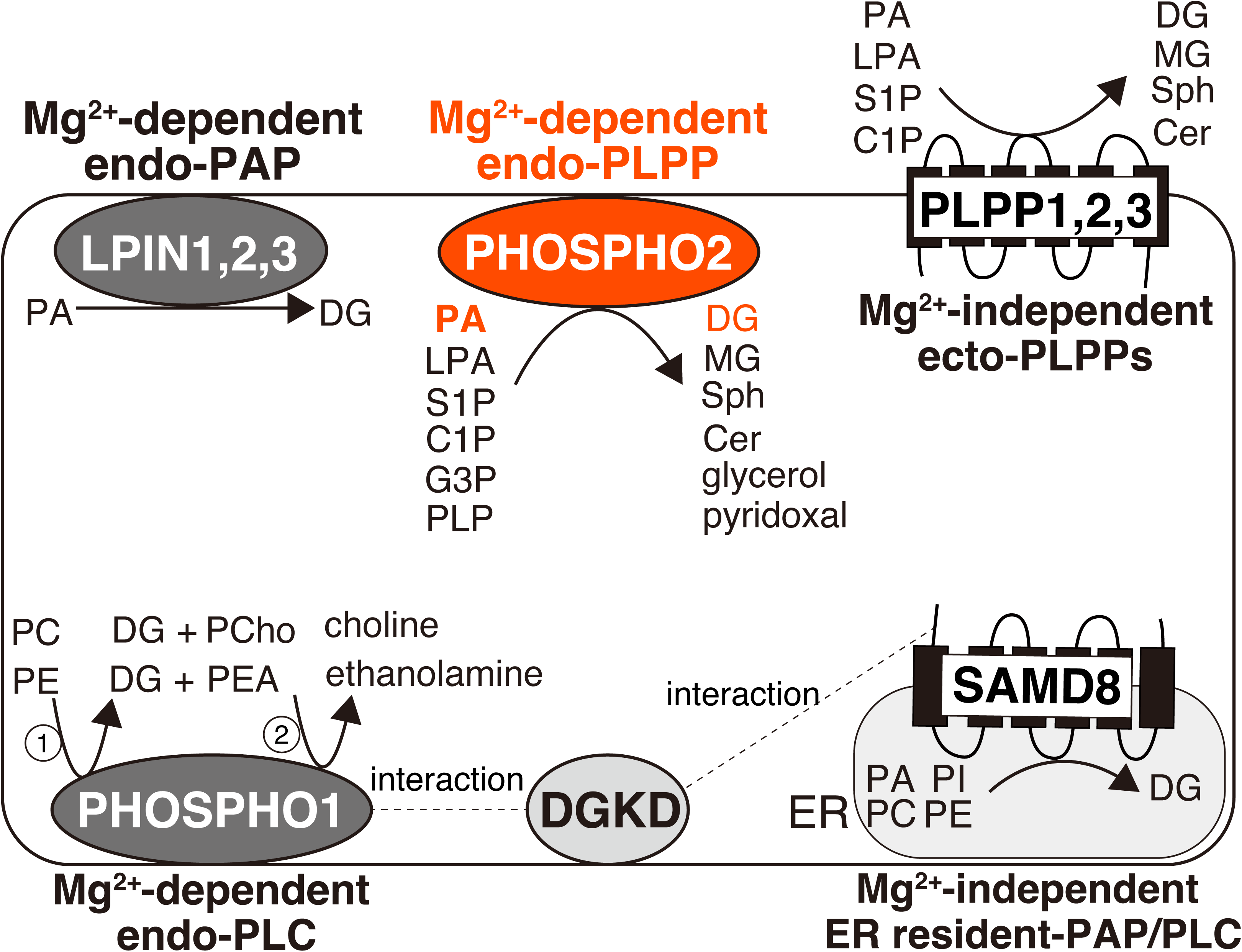
Schematic representation of the mammalian lipid metabolizing enzymes. PHOSPHO2 exhibits phospholipid phosphatase (PLPP) activity (EC 3.1.3.113) toward PA (EC 3.1.3.4), LPA (EC 3.1.3.106), S1P (EC 3.1.3.114), C1P (EC 3.1.3.115), and G3P phosphatase activity (EC 3.1.3.21) in addition to PLP phosphatase activity (EC 3.1.3.74). Because the PAP activity of PHOSPHO2 is activated in the presence of PC or PE (Fig. 5B), PHOSPHO2 may primarily hydrolyze PA on the cellular membrane. PLPPs (Gene names: PLPP1, 2, and 3) are Mg^2+^-independent ectoenzymes that hydrolyze phospholipids (PA, LPA, S1P and C1P) on the cellular surface. Lipin enzymes (Gene names: LPIN1, 2, and 3) are Mg^2+^-dependent cytosolic phosphatases that hydrolyze only PA [18]. PHOSPHO1 (42% sequence identity with PHOSPHO2) exhibits PC-PLC (EC 3.1.4.3) and PE-PLC (EC 3.1.4.62), and PCho/PEA phosphatase (EC3.1.3.75) activities [38]. Sphingomyelin synthase-related protein (SMSr) (Gene name: SAMD8, NCBI Gene ID:142891, UniProt accession number: Q96LT4), an ER-resident transmembrane protein containing active site motifs common to PLPPs [86], exhibits PAP and PLC activities toward PC, PE, and PI [43]. Unlike PHOSPHO2, PHOSPHO1 and SMSr preferably hydrolyze SFA- and/or MUFA-containing glycerophospholipids compared to PUFA-containing phospholipids [43]. Moreover, SMSr and PHOSPHO1 interact with DGKδ isozyme (Gene name: DGKD) [38, 40, 63], which preferably phosphorylates SFA- and/or MUFA-containing DGs [74]. However, interaction between PHOSPHO2 and DGKδ was not observed [38]. In contrast to other DG-generating enzymes, PHOSPHO2 localizes to the cytosolic fraction and displays Mg^2+^-dependent PLPP activity.

We recently identified human PLC or PAP enzymes that selectively produce SFA- and/or monounsaturated fatty acid-containing DG species such as sphingomyelin synthase (SMS)-related protein (SMSr) (Gene name: SAMD8, NCBI Gene ID:142891, UniProt accession number: Q96LT4) [43, 63], SMS isoform 1 (SMS1) (Gene name: SGMS1, NCBI Gene ID: 259230, UniProt accession number: Q86VZ5) [39], SMS isoform 2 (SMS2) (Gene name: SGMS2, NCBI Gene ID: 166929, UniProt accession number: Q8NHU3) [41], and PHOSPHO1[38]. SMS proteins (SMS1, SMS2, and SMSr) are transmembrane proteins in which the active-site motifs of PLPPs are highly conserved [86]. SMS1 and SMS2 act as PC-PLC and PE-PLC in addition to SM synthesis (EC 2.7.8.27). SMSr has no SMS activity, but displays multi-glycerophospholipid PLC hydrolase activity, including PAP, PI-PLC, PC-PLC, and PE-PLC. Because these proteins and PHOSPHO1 preferentially generate SFA- and/or MUFA-containing DG, these enzymes may be involved in an alternative DG signaling pathway independent of PI turnover (PUFA-containing DG signaling pathway) [74]. Moreover, the intracellular localizations of these new enzymes are distinct (Fig. 14). We found that DGKδ interacted with SMSr and PHOSPHO1 [38, 63], and DGK ζ isozyme (Gene name: DGKZ, NCBI Gene ID: 8525, UniProt accession number: Q13574) interacted with SMS1 and SMSr [40] (Fig. 14). Notably, both DGKδ and DGKζ phosphorylate SFA and/or MUFA-containing DG species which are not derived from the PI cycle [74]. Taken together, we assume that there are alternative DG signaling pathways involving new SFA-DG– generating enzymes, DGKδ and DGKζ that are independent of PI turnover. As PHOSPHO2 showed higher PAP activity toward medium-chain fatty acid-containing PA and PUFA-containing PA, PHOSPHO2 may be involved in new DG signaling distinct from SFA and/or MUFA-containing DG signaling.

An important limitation of this study is that our findings are based on *in vitro* experiments and *in silico* analyses of PHOSPHO2. Although the biological significance of PHOSPHO2 *in vivo* remains poorly understood, our findings provide evidence that PHOSPHO2 can act as a PLPP under the conditions we examined. These findings provide a basis for further investigation into the molecular mechanisms underlying the physiological functions of PHOSPHO2 *in vivo*. The biological role of PHOSPHO2 in lipid metabolism *in vivo* remains an interesting topic for future investigation.

In summary, we showed that human PHOSPHO2 displayed Mg^2+^-dependent PLPP activity *in vitro*. Unlike the Lipin family (PAP1), PHOSPHO2 is able to break down S1P, C1P, and LPA in addition to PA. Moreover, in contrast to the conventional PLPP family (PAP2), PHOSPHO2 can hydrolyze PA, S1P, C1P, and LPA on the membranes facing the cytoplasm. Both PHOSPHO1 (PC/PE-PLC) and PHOSPHO2 (PAP1/PLPP) can hydrolyze phospholipids, but their substrate specificities are distinct. Hence, PHOSPHO2 can be categorized as a novel mammalian PLPP (Mg^2+^-dependent endo-PLPP), which is distinct from conventional PLPPs (Mg^2+^-independent ecto-PLPP), Lipin (Mg^2+^-dependent PAP1), and PLCs.

## Experimental procedures

### Materials

Lipids: 1-pentadecanoyl-2-oleoyl-*sn*-glycerol (15:0/18:1-DG, Cat. No. 330722), 1-palmitoyl-2-oleoyl-*sn*-glycero-3-phosphate (16:0/18:1-PA, Cat. No. 840857), 1-palmitoyl-2-oleoyl-*sn*-glycero-3-phosphoinositol (16:0/18:1-PI, Cat. No. 850142), 1-palmitoyl-2-oleoyl-*sn*-glycero-3-phospho-L-serine (16:0/18:1-PS, Cat. No. 840034), 1-palmitoyl-2-oleoyl-*sn*-glycero-3-phosphocholine (16:0/18:1-PC, Cat. No. 850457); 1-palmitoyl-2-oleoyl-*sn*-glycero-*sn*-glycerol (16:0/18:1-DG, Cat. No. 800815), 1-palmitoyl-2-oleoyl-*sn*-3-phosphoglycerol (16:0/18:1-PG, Cat. No. 840457), 1-stearoyl-2-hydroxy-sn-glycero-3-phosphate (18:0-LPA, Cat. No. 857128), 1-myristoyl-2-hydroxy-sn-glycero-3-phosphate (14:0-LPA, Cat. No. 857120), D-erythro-sphingosine-1-phosphate (d20:1-S1P, Cat. No. 860662), 1,2-dilauroyl-sn-glycero-3-phosphate (12:0/12:0-PA, Cat. No. 840635) 1,2-distearoyl-sn-glycero-3-phosphate (18:0/18:0-PA, Cat. No. 830865), 1,2-dipalmitoyl-sn-glycero-3-phosphate (16:0/16:0-PA, Cat. No. 830855), 1-stearoyl-2-arachidonoyl-sn-glycero-3-phosphate (18:0/20:4-PA, Cat. No. 840863), and 1,2-dioleoyl-sn-glycero-3-phosphate (18:1/18:1-PA, Cat. No. 840875) were purchased from Avanti Polar Lipids (Alabaster, AL, USA). 1-palmitoyl-2-hydroxy-sn-glycero-3-phosphate (16:0-LPA, Cat. No. 10010290), 1-oleoyl-2-hydroxy-sn-glycero-3-phosphate (18:1-LPA, Cat. No. 62215), 1-arachidonoyl-2-hydroxy-sn-glycero-3-phosphate (20:4-LPA, Cat. No. 10019), D-erythro-sphingosine-1-phosphate (d18:1-S1P, d17:1-S1P, and d16:1-S1P) (Cat. No. 62570, 22498, and 9002921, respectively). N-octanoyl-D-erythro-sphingosine-1-phosphate (d18:1/8:0-C1P, Cat. No. 62547), N-stearoyl-D-erythro-sphingosine-1-phosphate (d18:1/18:0-C1P, Cat. No. 39228), D-erythro-dihydrosphingosine (sphinganine (d18:0), Cat. No. 10007945), and D-erythro-sphingosine (Sph, Cat. No. 10007907) were purchased from Cayman Chemical Company (Ann Arbor, MI, USA).

Antibodies: mouse monoclonal anti-Strep II antibody (M211-3) was purchased from Medical and Biological Laboratories (Nagoya, Japan). Peroxidase-conjugated goat anti-mouse IgG was purchased from Bethyl Laboratories (Montgomery, TX, USA). Peroxidase-conjugated goat anti-rabbit IgG antibody (111-036-045) was purchased from Jackson ImmunoResearch (West Grove, PA, USA).

Detergents: n-dodecyl-β-D-maltoside (DDM) was purchased from Cayman Chemical Company. Cholesteryl hemisuccinate (CHS) was purchased from Sigma‒Aldrich (St. Louis, MO, USA).

Compounds: D-Desthiobiotin (Cat. No. D1411) and (±)-Propranolol hydrochloride (Cat. No. P0884) were purchased from Sigma‒Aldrich. Linear polyethylenimine hydrochloride (transfection grade, MW 40,000) was purchased from Polysciences (Warrington, PA, USA). Pyridoxal Phosphate Monohydrate (Cat. No. 165-20943), Isopropyl-β-D(-)-thiogalactopyranoside (IPTG) (Cat. No. 094-05144), and N-ethylmaleimide (Cat. No. 058-02061) were purchased from Wako Pure Chemicals (Osaka, Japan).

### Plasmids

First, a C-terminally Twin-Strep-tagged human phosphatase orphan 2 (PHOSPHO2) (NCBI Reference Sequence: NM_001008489.4, UniProt accession number: Q8TCD6) expression plasmid was generated (Addgene plasmid # 222016; RRID:Addgene_222016) [39–41]. Briefly, we used pCAGGS-N-TEV-Twin-Strep (Addgene plasmid # 202526; http://n2t.net/addgene:202526; RRID: Addgene_202526), a plasmid encoding a C-terminally TEV (Tobacco Etch Virus protease)-cleavable Twin-Strep tag (ENLYFQGS-WSHPQFEK-(GGGS)_2_-GGSA-WSHPQFEK) fusion protein in mammalian cells. The plasmid was linearized using inverse PCR with the following primers: forward, 5′-CTCGAGGAAAACCTCTATTTTCAAGGC -3′; reverse, 5′-CATGGTGGCTCGAGCGGCCGCGATATC -3′. Full-length PHOSPHO2 coding region was amplified from cDNA isolated from HEK293 cells using polymerase chain reaction (PCR) using the following primers: forward, 5′-GCTCGAGCCACCATGAAAATTTTGCTAGTTTTTG -3′; reverse, 5′-AGAGGTTTTCCTCGAGATCCTTTATTAGAAATTGTAAATGAGAAATTATATC -3′.

The amplified DNA was subcloned into the linearized pCAGGS-N-TEV-Twin-Strep vector using In-Fusion cloning (Clontech-Takara Bio). To express PHOSPHO2-TS in *E. coli*, pGEX-PHOSPHO2-TEV-Twin-Strep vector (Addgene plasmid # 222017; RRID:Addgene_222017) was generated. Briefly, the amplified DNA was inserted into pGEX-N-TEV-Twin-Strep vector (Addgene plasmid # 222014; RRID: Addgene_ 222014), a plasmid encoding C-terminally TEV protease-cleavable Twin-Strep tag in *E. coli*, via In-Fusion cloning.

### Mammalian cell culture and expression of PHOSPHO2 proteins

HEK293 cells (Japanese Collection of Research Bioresources, Tokyo, Japan) were maintained in Dulbecco’s modified Eagle medium (DMEM, Wako Pure Chemicals, Osaka, Japan) supplemented with 5% fetal bovine serum (FBS, Thermo Fisher Scientific, Waltham, MA, USA) and 100 U/mL penicillin/100 μg/mL streptomycin (Wako Pure Chemicals) at 37°C in an atmosphere containing 5% CO_2_. HEK293 cells (6.25 × 10^5^) were plated on 150-mm dishes. After 96 h, the cells were transiently transfected with the plasmids using polyethylenimine (PEI) Max (#24765-100, Polysciences, Warrington, PA, USA). Plasmid DNA and PEI (1 mg/mL, pH 8.0) were preincubated for 10 min at a 1:3 ratio (15 µg DNA: 45 µL PEI) in 750 µL of Opti-MEM (Cat. No. 31985-070, Gibco, Waltham, MA, USA) before transfection. The DNA-PEI complex (15 µg DNA) was transfected into HEK293 cells (approximately 50% confluent cells in a 150-mm dish). After 24 h, the cells were harvested using a scraper and centrifugation. The cell pellets were resuspended in phosphate-buffered saline (PBS) containing 40% (v/v) glycerol. The cell samples were then flash-frozen in liquid nitrogen and stored at –80°C for further use.

### Expression of PHOSPHO2-TS proteins in E. coli

*E. coli* BL21 cells (Cat. No. 9126, Takara Bio, Shiga, Japan) harboring the expression plasmid (pGEX-PHOSPHO2-TEV-Twin-Strep) were grown at 37°C in LB medium supplemented with 100 μg/mL ampicillin. The growth of the cells was determined by measuring the optical density at 600 nm (OD600). To induce protein expression, 0.5 mM IPTG was added to the culture when OD600 reached 0.5. After incubation at 37°C for 3 h, cells were harvested using centrifugation, and pellets were resuspended in PBS containing 40% (v/v) glycerol. The cell samples were flash-frozen in liquid nitrogen and stored at –80°C for further use.

### Purification of Twin-Strep-tagged proteins

Human PHOSPHO2-TS expressing cells (HEK293, cell pellets from 32 × 150-mm dishes; BL21 cells, cell pellets from 4 liters of culture medium) were thawed at 4°C. After centrifugation at 10,000 × g for 20 min at 4°C, cell pellets were resuspended in ice-cold lysis buffer (20 mM Tris-HCl (pH 7.4), 2 mM MgCl_2_, 150 mM NaCl, 0.1 mM DTT, 10% (v/v) glycerol, 1 mM phenylmethylsulfonyl fluoride (PMSF), 20 mg/mL aprotinin, 20 mg/mL leupeptin, and 20 mg/mL pepstatin) and cells were homogenized on ice using a homogenizer (Cat. No. 885300-0100, Kimble Kontes, Vineland, NJ, USA) and lysed using sonication on ice. The cell lysate was gently mixed by rotation for 20 min at 4°C. Insoluble material was removed by two rounds of centrifugation (10,000 × g, 20 min at 4°C). The protein was purified using Strep-Tactin XT beads (IBA Life Sciences, Goettingen, Germany) [39–41]. The supernatant was applied to a column containing 1,000 µL of Strep-Tactin XT resin. The beads were washed with wash buffer (20 mM Tris-HCl (pH 7.4), 2 mM MgCl_2_, 150 mM NaCl, 0.1 mM DTT, and 10% (v/v) glycerol). Twin-Strep-tagged proteins were eluted with elution buffer (wash buffer containing 2 mM D-desthiobiotin). Purified samples were concentrated using Amicon Ultra-15 centrifugal filter devices (10 kDa molecular weight cutoff, Cat. No. UFC901024, Merck, Darmstadt, Germany).

### Sodium dodecyl sulfate-polyacrylamide gel electrophoresis (SDS-PAGE) and Immunoblot analysis

The purified proteins were mixed with 5 × Laemmli sample buffer containing 2.5% (v/v) 2-mercaptoethanol and incubated at 95°C for 10 min. The samples were analyzed using SDS-PAGE. Twin-Strep-tagged proteins were detected using immunoblot analysis as previously described [38, 41]. BlueStar Prestained Protein Ladder (Cat. No. #RPN2106, Nippon Genetics, Tokyo, Japan) was used as a molecular mass marker. Immune complexes were visualized with Amersham ECL Western Blotting Detection Reagent (Cytiva, Marlborough, Massachusetts, USA) and LuminoGraph I (WSE-6100, ATTO, Tokyo, Japan). Proteins were also visualized using Coomassie Brilliant Blue staining with Coomassie Brilliant Blue staining [87], and the stained gels were scanned using a flatbed scanner (GT-X980, Epson, Nagano, Japan). Adjustments to brightness and contrast were uniformly applied across the entire image.

### Preparation of DDM and CHS detergents stock solutions

To prepare the detergent stock solution (10% (w/v) DDM and 2% (w/v) CHS), 5 g of DDM was added to 40 mL of 200 mM Tris-HCl (pH 8.0), followed by gentle rotation until the DDM was completely dissolved. CHS (1 g) was added to the DDM solution and sonicated until the solution became translucent. After sonication, 200 mM Tris-HCl (pH 8.0) was added up to 50 mL and incubated at 25°C with gentle rotation until the solution became transparent. The detergent solution was stored at –25°C.

### Preparation of micelles

The reaction solution for the phosphatase activity assay was prepared as previously described [38, 41]. Phospholipids dissolved in chloroform/methanol (2:1 (v/v)) were dried under a stream of N_2_ gas to form a lipid film on the vial wall. The lipid film was resuspended in the reaction buffer (20 mM Tris-HCl (pH 7.4), 2 mM MgCl_2_, 150 mM NaCl, 0.04% (w/v) DDM, 0.008% (w/v) CHS, 100 µM dithiothreitol, and 10% (v/v) glycerol) to a final lipid concentration of 400 µM each. The reaction solutions containing lipids were vortexed for 3 min and sonicated using a bath sonicator (Sonifier® Model 450, Branson, St. Louis, MO, USA) four times for 3 min each at 65°C.

### In vitro phosphatase activity assay

DG-generating activities (PAP/PLC activity) of the purified proteins were evaluated using previously reported methods [38, 41]. All enzyme activity assays were performed in a detergent-based micelle environment. The purified PHOSPHO2-TS samples (10 µL containing approximately 500 ng of protein purified from HEK293 cells or 4 µg of protein purified from *E. coli*) were diluted in 10 µL ice-cold reaction buffer on ice (final concentration of 20 mM Tris-HCl (pH 7.4), 150 mM NaCl, 0.05% (w/v) DDM, 0.01% (w/v) CHS, 0.1 mM dithiothreitol, 10% (v/v) glycerol, and 50 µM phospholipids (16:0/18:1-PC, 16:0/18:1-PE, 16:0/18:1-PA, 16:0/18:1-PI, 16:0/18:1-PS, or 16:0/18:1-PG). For the phosphatase activity assay, lipids (LPA, S1P, or C1P) were added to the reaction buffer at a final concentration of 200 µM. After incubation (enzyme reaction) for 60 min at 37°C, the assay samples were stored at –80°C until further analysis.

### DG quantification using LC–MS/MS

Lipids in the samples were extracted using the Bligh and Dyer method . Briefly, samples (20 µL) were added to 380 µL of water. To compensate for any possible variation during the entire process (*e.g.,* recovery rate of lipids in the lipid extraction step, HPLC injection, and ionization variability) , 20 ng of 15:0/18:1-DG was added as an internal standard (I.S.) from this step. The samples were mixed with methanol (1 mL) and chloroform (0.5 mL). After vortexing for 30 s, samples were incubated for 5 min at room temperature (25°C). The samples were then mixed with chloroform (0.5 mL) and water (0.5 mL) and vortexed for 30 s. After centrifugation (1,000 × g, 25°C, 10 min), the lower phase containing extracted lipids was decanted to a new test tube and used for quantitation of DG via LC‒ MS/MS [38, 41, 42]. Ionized DG molecular species ([M + NH4]^+^) were isolated in the first quadrupole (Q1). Thereafter, the product ions of the DG species were reselected at Q3 after fragmentation at Q2 by collision-induced dissociation. Peak regions corresponding to analyte signals in chromatograms were selected. To calculate the peak area of analytes, peak areas for each MRM transition were integrated using MultiQuant (Version 3.0.3) (AB SCIEX, Framingham, MA). The amounts of detected lipids were normalized by calculating the peak area ratio of the analyte to the internal standard (Peak Area Ratio) for relative quantification. For absolute quantification of lipids, we generated calibration curves of Peak Area Ratio using commercially available purified lipids (16:0/18:1-DG) [41]. Using the calibration curves, we determined the absolute amounts of 16:0/18:1-DG in samples (mole/sample).

### Phosphate quantification

Phosphatase activity of PHOSPHO2 was measured using the BIOMOL Green reagent (Enzo Life Sciences, Farmingdale, USA). Samples (5 µL) and 45 µL of the wash buffer (20 mM Tris-HCl (pH 7.4), 2 mM MgCl_2_, 150 mM NaCl, 0.1 mM DTT, and 10% (v/v) glycerol) were added to 96-well clear microplates (Cat. No. 208004, Porvair Sciences, Wrexham, UK). The BIOMOL Green reagent (100 µL) was added, and the plates were incubated for 30 min at 25°C. The absorbance at 600 nm of each well was measured. For absolute quantification of free phosphate, calibration curves were generated using KH_2_PO_4_ at concentrations of 0, 0.63, 1.25, 2.5, 5, 10, 20, and 40 µM (50 µL/well).

### Protein alignment, structure prediction, and visualization

Multiple sequence alignment was created using ClustalW [88]. Structure predictions of human PHOSPHO2 and PHOSPHO1 containing a single Mg^2+^ ion were performed using AlphaFold3 [46]. The chemical structures of phosphorus compounds were drawn with the Chemical Sketch Tool from the Protein Data Bank website (https://www.rcsb.org/chemical-sketch). The putative substrates were docked into AlphaFold3-generated PHOSPHO1 and 2 models using SwissDock [48]. Three independent docking runs were conducted for each substrate-enzyme complex, and the binding mode with the lowest binding energy (ΔG (kcal/mol)) in which the phosphorus atom (P) of the docked substrate was located within 6 Å of the Mg^2+^ ion in PHOSPHO2 was selected for further analysis. The molecular docking results were visualized using UCSF Chimera (version 1.19) [89] and PyMOL (open-source version 3.1.0, https://pymol.org/). The substrate-binding pockets of Mg^2+^-binding PHOSPHO1 and PHOSPHO2 were predicted using the Computed Atlas of Surface Topography of the universe of protein Folds (CASTpFold) server [47]. The volumes of the putative substrate-binding pockets were calculated using CASTpFold. The volumes of substrates were calculated using MoloVol [90].

### Statistical analysis

Data are presented as means ± standard deviation (S.D.) and were analyzed using Student’s *t* test (two-tailed, unpaired) for the comparison of two groups or one-way analysis of variance (ANOVA) followed by Tukey’s post hoc test (for comparing means) or Dunnett’s post hoc test (for comparing each mean to a control mean) using GraphPad Prism 9 (GraphPad Software, Boston, MA, USA) to determine any significant differences. Individual data points (technical replicates) are superimposed on bar graphs. Statistical significance was set at *p* < 0.05.

## Data availability statement

Data supporting the findings of this study are available from the corresponding author upon request.

## Supporting information

This article contains supporting information.

## CRediT authorship contribution statement

**Kyoko Atsuta-Tsunoda**: Investigation, Writing – review & editing. **Chiaki Murakami**: Conceptualization, Data curation, Formal analysis, Funding acquisition, Investigation, Methodology, Project administration, Resources, Supervision, Validation, Visualization, Writing – original draft, Writing – review & editing. **Hiromichi Sakai**: Writing – review & editing. **Fumio Sakane**: Funding acquisition, Resources, Supervision, Writing – review & editing.

## Acknowledgements

We thank Mr. Takuma Kawai, Mr. Tomoki Hirai, Mr. Yasuhisa Hijikata, and Mr. Sho Inomata for technical assistance.

## Abbreviations

Cer: ceramide
C1P: ceramide-1-phosphate
DG: diacylglycerol
DGK: diacylglycerol kinase
G2P: glycerol-2-phosphate (β-glycerophosphate)
G2P: glycerol-2-phosphate (β-glycerophosphate)
G3P: glycerol-3-phosphate (α-glycerophosphate)
GP: glycerophosphate
HAD: haloacid dehalogenase
IPTG: Isopropyl-β-D(–)-thiogalactopyranoside
LC: liquid chromatography
LPA: lysophosphatidic acid
LPP: lipid phosphate phosphatase
MG: monoacylglycerol
MS: mass spectrometry
MUFA: monounsaturated fatty acids
NEM: N-Ethylmaleimide
PA: phosphatidic acid
PAP: PA phosphatase
PC: phosphatidylcholine
PE: phosphatidylethanolamine
PG: phosphatidylglycerol
Pi: inorganic phosphate
PI: phosphatidylinositol
PLC: phospholipase C
PLD: phospholipase D
PLP: Pyridoxal phosphate
PLPP: phospholipid phosphatase
PS: phosphatidylserine
S1P: sphingosine-1-phosphate
SFA: saturated fatty acids
Sph: sphingosine
PUFA: polyunsaturated fatty acids
TEV: Tobacco Etch Virus
PEI: polyethylenimine
PMSF: phenylmethylsulfonyl fluoride
TS: Twin-Strep-tag.

---

**Suppl. Figure S1.**
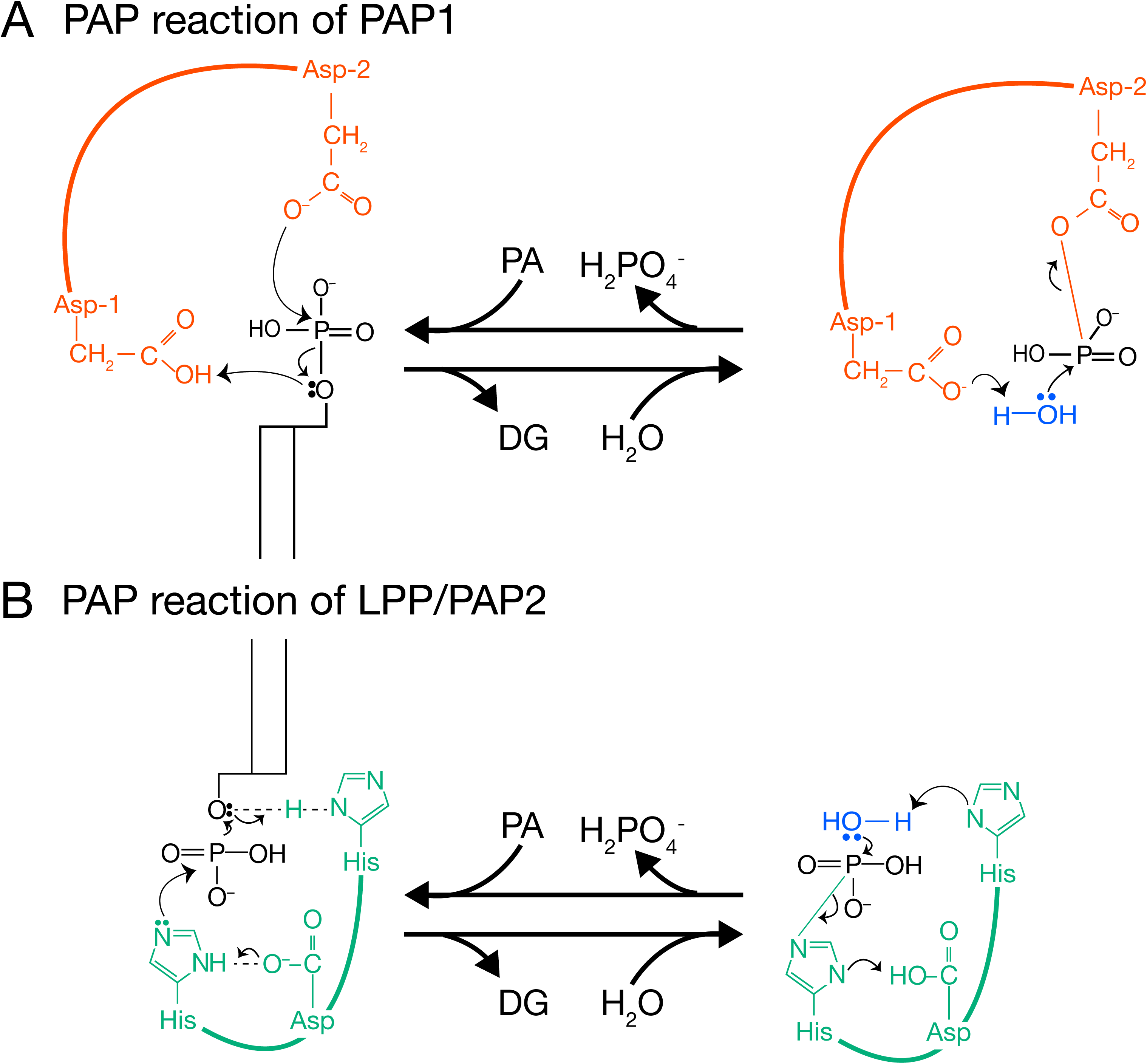
Proposed catalytic mechanism for PAP1 and PAP2. (A) Mg^2+^-dependent PAP (PAP1) proceeds in two steps [9]. In the first step, the carboxylate group of the catalytic aspartate (Asp-1) acts as the nucleophile and forms an aspartyl phosphate intermediate, while the cleaved group pulls a hydrogen from the second Asp (Asp-2) to form DG. In the second step, a water molecule activated by the second Asp attacks the aspartyl-phosphate intermediate, and free phosphate is generated. (B) The reaction of PLPP (PAP2/LPP) involves a charge relay system [12]. The first step serves to conduct electrons through hydrogen bonds from the catalytic Asp to His, thereby forming a bond between His and the polar head of phospholipid (N–P bond). The catalytic histidine facilitates the cleavage of the phosphodiester bond of phospholipids and release of DG by acting as a general acid. In the second step, His acts as a general base to facilitate the cleavage of the N–P bond. The water molecule serves as a general acid.

**Suppl. Figure S2.**
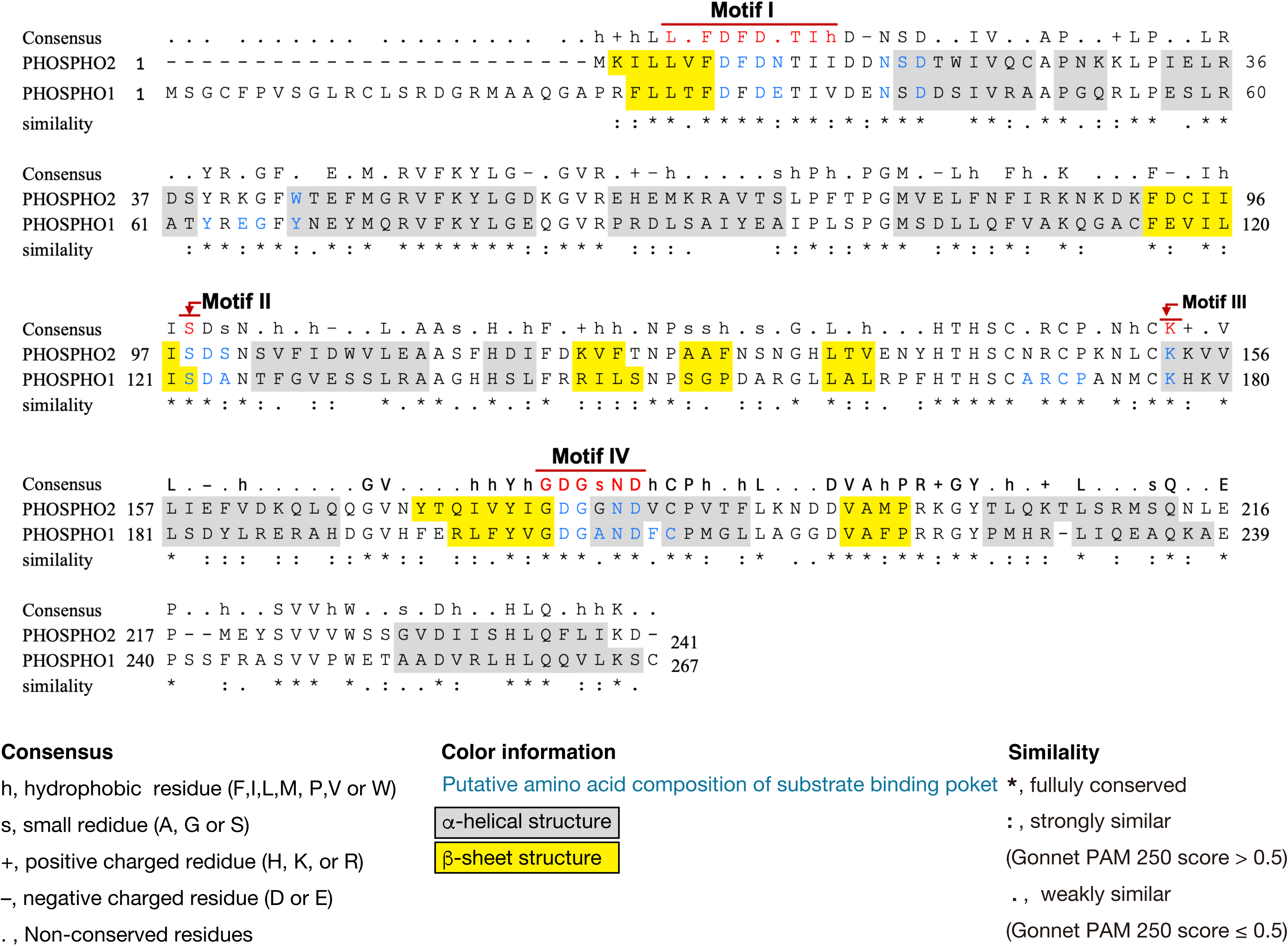
Sequence alignment of human PHOSPHO2 and PHOSPHO1. Sequence alignment of the human PHOSPHO1 (NCBI Reference Sequence: NP_848595.1, UniProt accession number: Q8TCT1) and PHOSPHO2 (NCBI Reference Sequence: NP_001008489.1, UniProt accession number: Q8TCD6) was created using ClustalW (version 2.1) provided from the DNA Data Bank of Japan (DDBJ) [88]. α-helix and β-sheet structures are highlighted in gray and yellow, respectively. Amino acid residues comprising the putative substrate binding pocket (Fig. 11D and Table 3) are colored in blue.

**Suppl. Figure S3.**
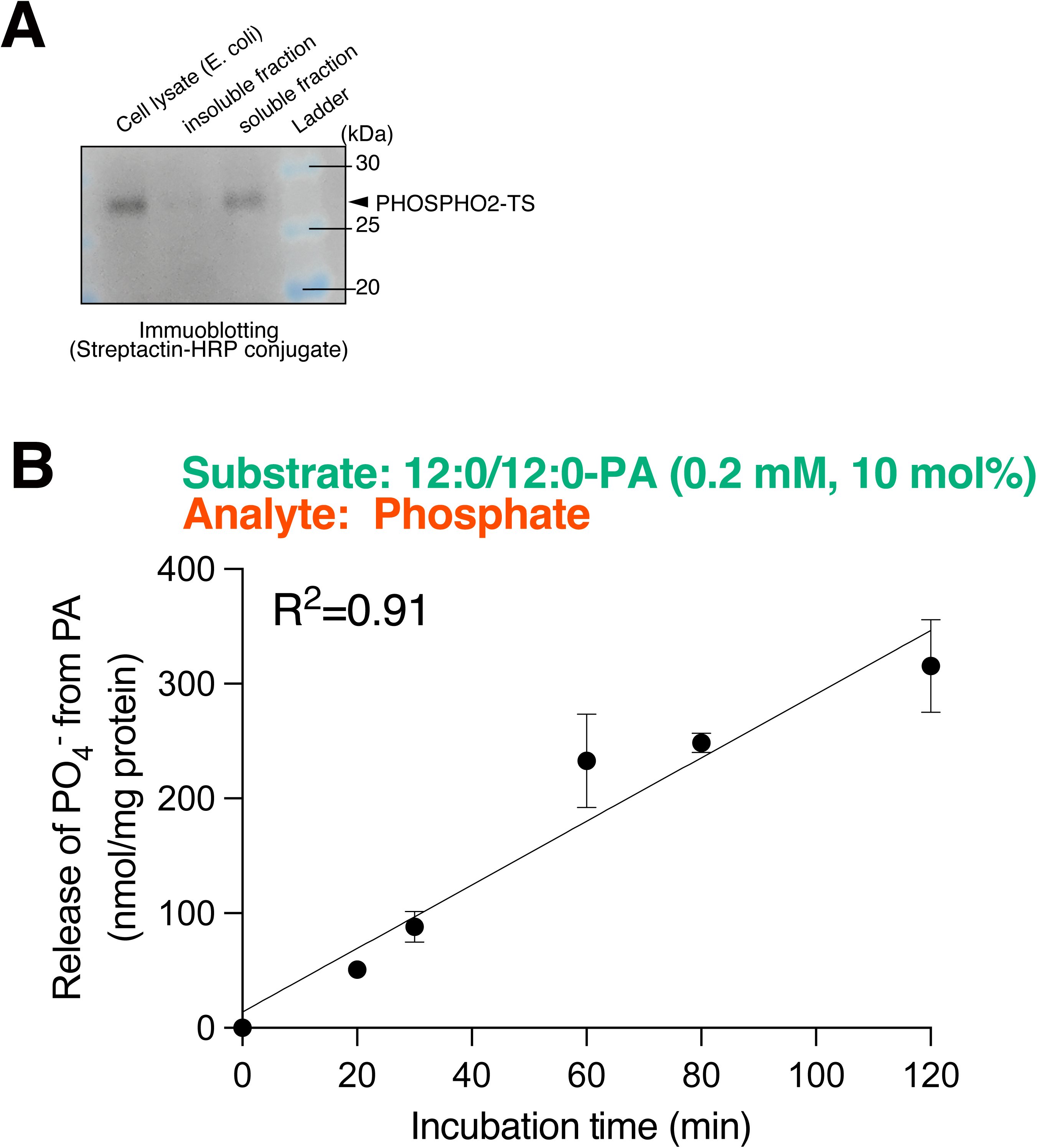
Validation of enzyme purification and activity assay. (A) Immunoblot analysis of the solubility of PHOSPHO2-TS expressed in *E. coli* (BL21). Cell lysates were separated into 10,000 *g* supernatant and pellets (insoluble fraction) and subjected to SDS-PAGE (12%) followed by immunoblot analysis using precision Protein Streptactin-HRP conjugate (#1610380, Bio-Rad, Hercules, California) Protein Ladder One Plus (Cat. No. #19593-25, Nacalai Tesque, Kyoto, Japan) was used as a molecular mass marker. (B) Dependence of PAP activity of PHOSPHO2 on time using 12:0/12:0-PA-containing mixed micelles. The release of P_i_ from 12:0/12:0-PA was measured using BIOMOL Green reagent. The surface and bulk concentrations of PA were 10 mol% and 0.2 mM, respectively. The enzyme activity with 4 µg PHOSPHO2 purified from *E. coli* (BL21) was measured for the indicated time intervals (0, 20, 30, 60, 80, and 120 min).

**Suppl. Figure S4.**
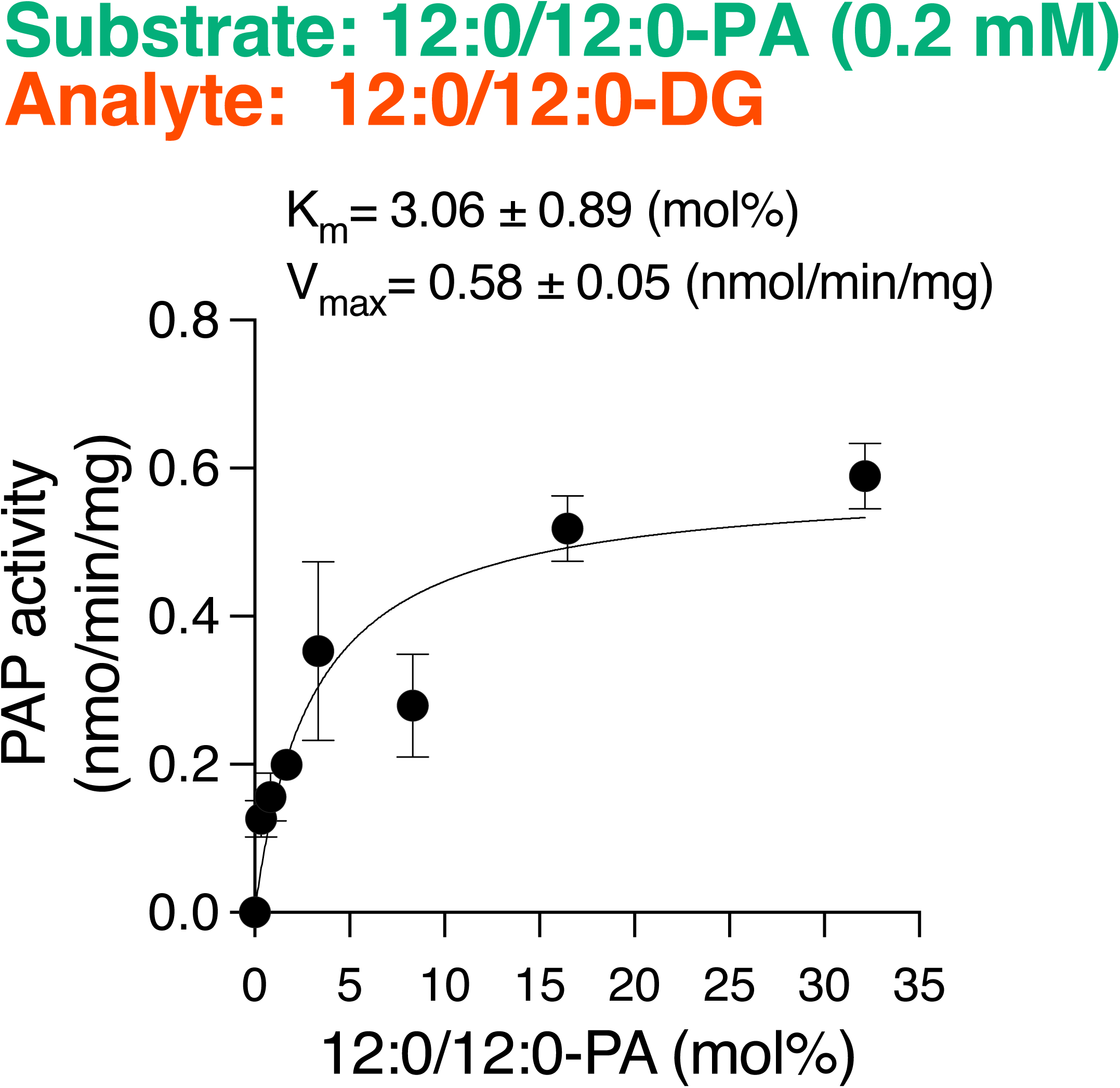
Kinetic analysis of PAP activities of PHOSPHO2. PAP activity was measured as a function of the indicated surface concentration (mol%) [45] using 12:0/12:0-PA-DDM/CHS mixed micelles. To evaluate phosphatase activities using phospholipids/DDM/CHS-mixed micelles, a surface dilution kinetic scheme was used. The mol% of lipids in the DDM/CHS/lipid mixed micelles were calculated using the formula: mol% = 100 × [lipid (mol)]/([DDM (mol)] + [CHS (mol)] + [lipid (mol)]). In the experiment, molar concentration of 12:0/12:0-PA was held constant at 200 µM, and the DDM/CHS concentration (5:1 ratio) was varied to obtain the indicated surface concentrations of 12:0/12:0-PA. Values represent averages of triplicate measurements. Data are presented as means ± S.D. (*n* = 3, technical replicates). The enzyme kinetic parameters (*K*_m_ and *V*_max_) were obtained by fitting kinetic models to the experimental data using GraphPad Prism 9.

**Suppl. Figure S5.**
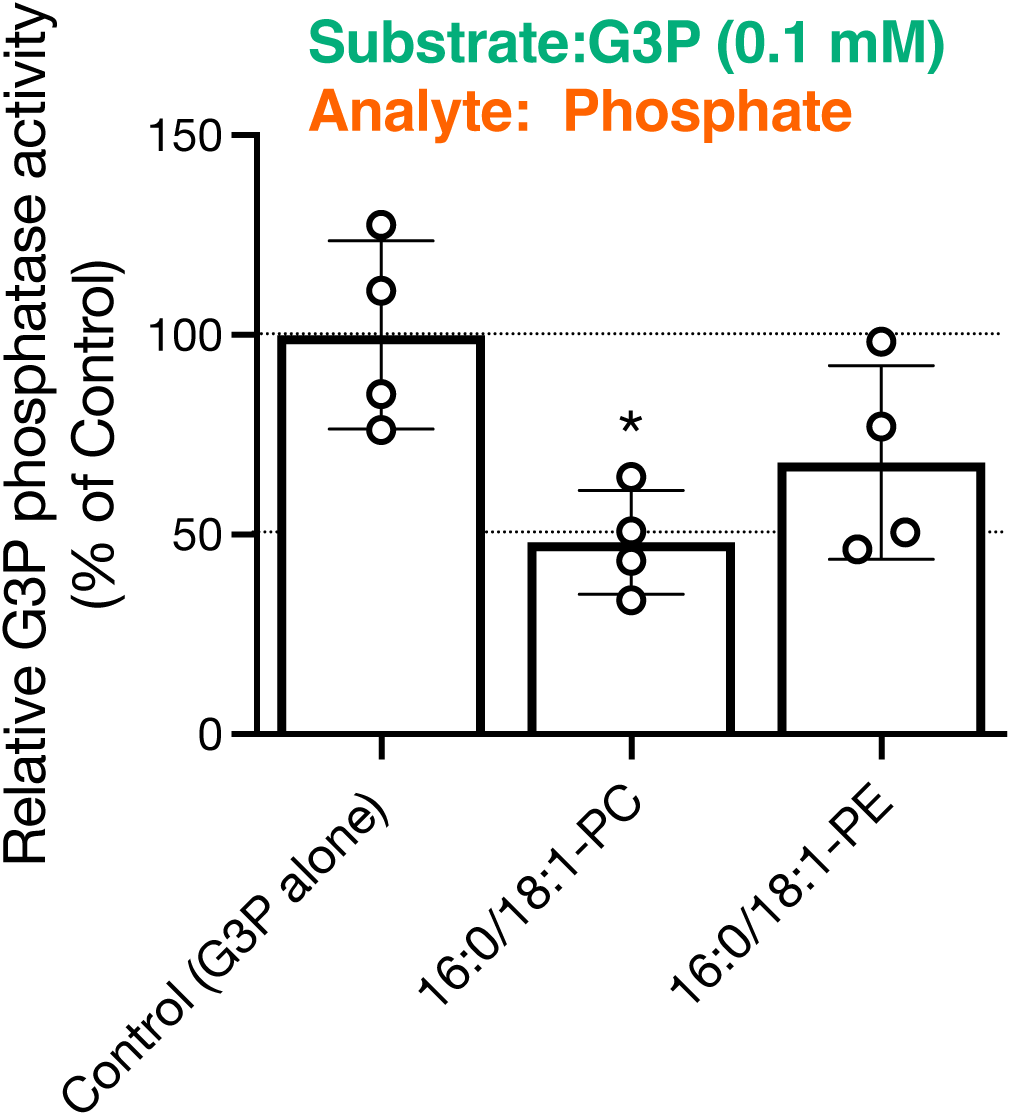
Effects of PC and PE on the G3P phosphatase activities of PHOSPHO2. G3P phosphatase activity of PHOSPHO2 was measured in the presence of PC or PE (each at 0.1 mM) in addition to G3P (0.1 mM). Values are presented as percentages of the G3P phosphatase activity of PHOSPHO2 in the presence of the substrate alone (set at 100%). Values are presented as the mean ± S.D. (*n* = 4, technical replicates). *, *p* < 0.05 (*vs.* control). One-way ANOVA with Dunnett’s post hoc test was used.

**Suppl. Figure S6.**
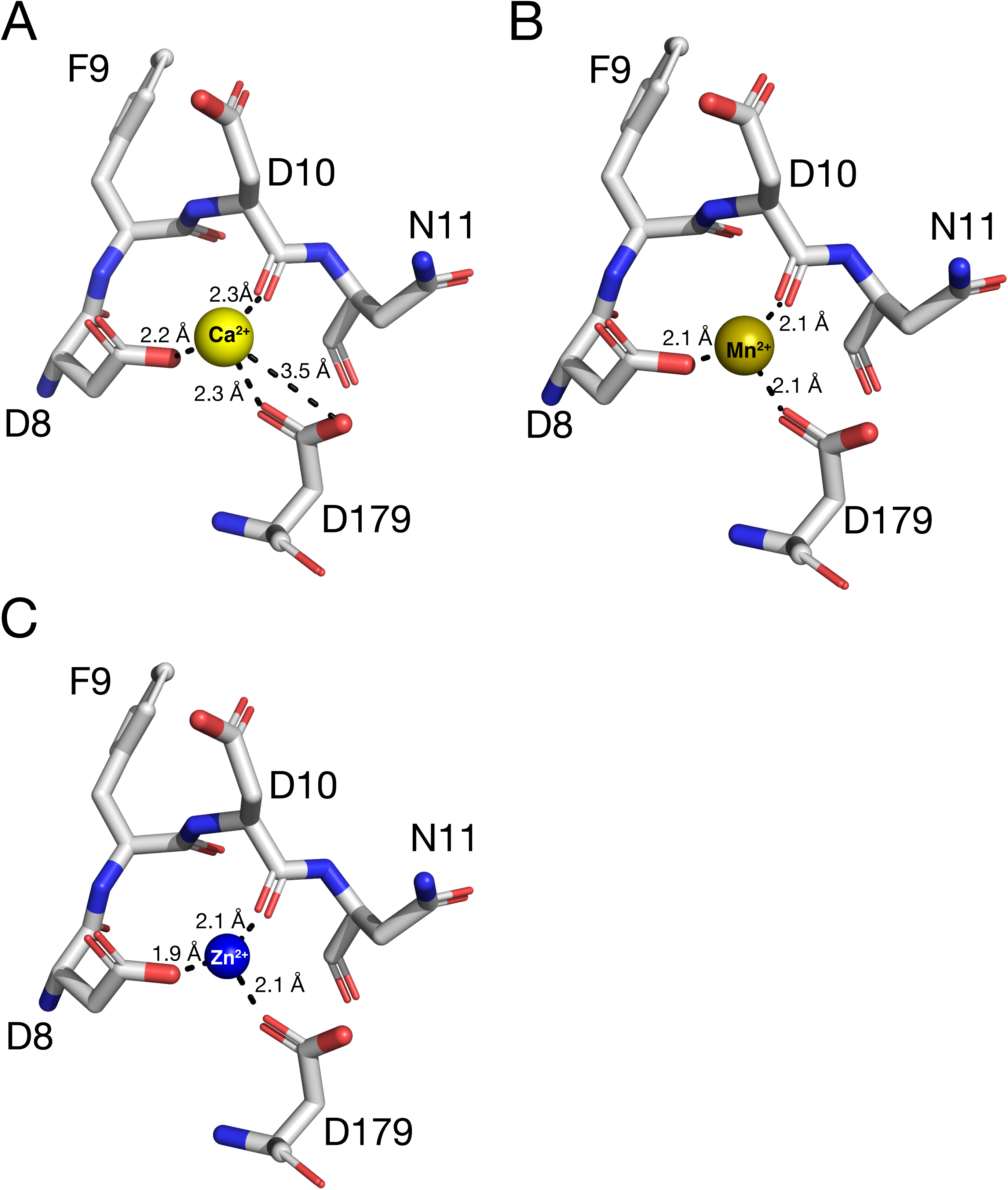
Active site of PHOSPHO2 bound to divalent metal ion. Predicted structures of human PHOSPHO2 bound to Ca^2+^ (A), Mn^2+^ (B), or Zn^2+^ (C) were generated using AlphaFold3 [46]. Contacts between metal ions and PHOSPHO2 were analyzed using UCSF ChimeraX (version 1.10) [89] and visualized using PyMOL. Dotted lines indicate coordination bonds. Carbon (grey), nitrogen (blue), and oxygen (red) are highlighted. The metal ions are represented as colored spheres.

**Suppl. Figure S7.**
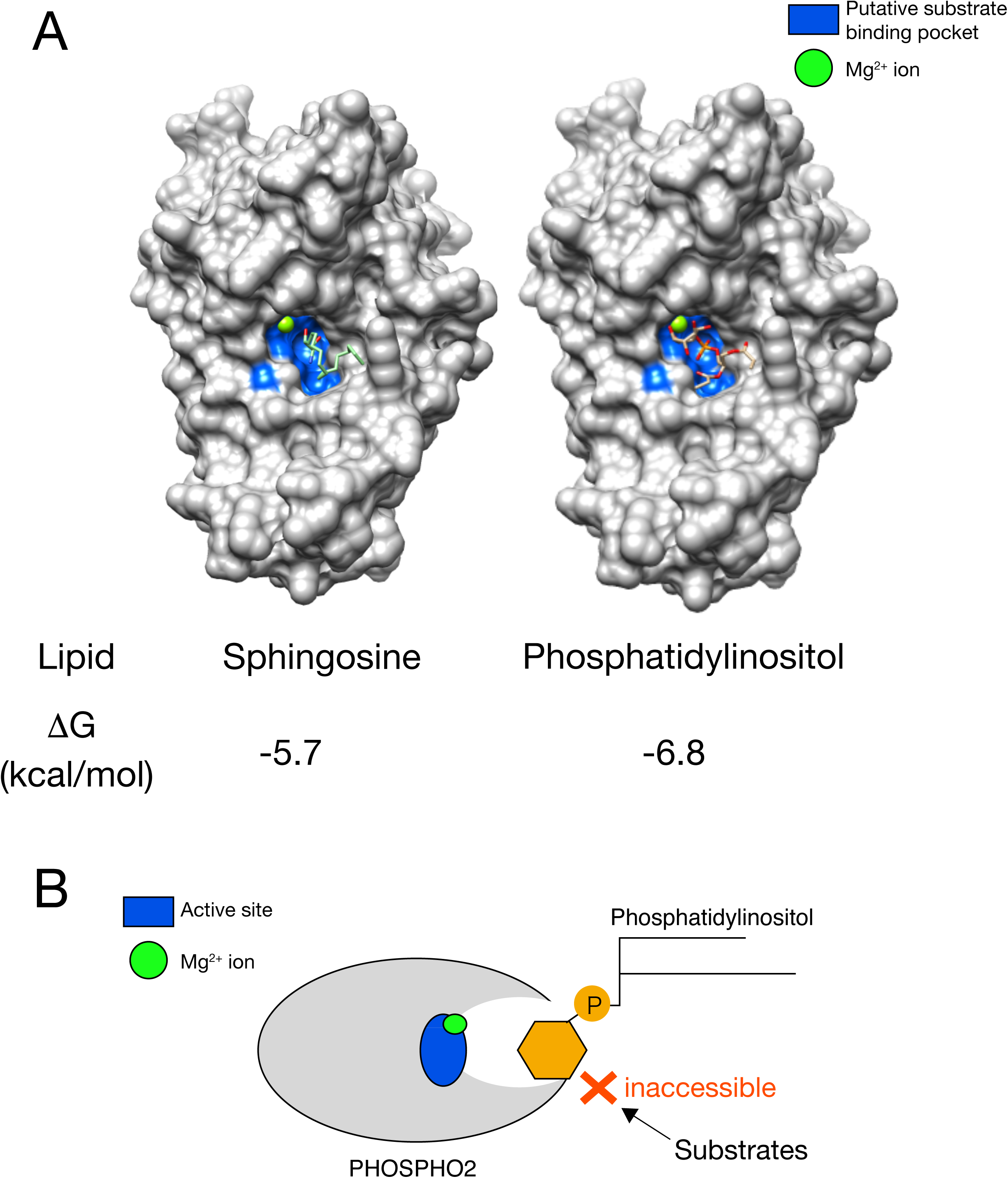
Docking simulations of PI and Sph to the AlphaFold3-predicted PHOSPHO2. (A) The docked model of PHOSPHO2 with Mg^2+^ interacting with *D*-erythro-Sphingosine (Sph) and 3:0/3:0-PI. The docking model was generated using Webina [49]. A model was visualized and analyzed using UCSF Chimera. (B) A model for the interaction of PHOSPHO2 with PI. Sph and PI may inhibit the phosphatase activity because they sterically hinder substrate access to the catalytic pocket.

**Suppl. Table S1.** Estimation of cellular concentrations of substrates for PHOSPHO2.

| Substrate | Estimated intracellular concentration ($\mu\text{M}$) | EC number [6] | References | Calculated intracellular enzyme activity of PHOSPHO2 (nmol/mg/min) | |
| --- | --- | --- | --- | --- | --- |
|  |  |  |  | In cytosol (in the absence of PC) | On the cellular membrane (PC-rich environment) |
| PLP | 3.5 | EC 3.1.3.74 | [61, 62] | **0.2** | **0.2** |
| PA | 30–300 | EC 3.1.3.4 (PAP1)<br>EC 3.1.3.113 (PLPP) | [52, 59, 63–66] | 16:0/18:1-PA: **0.6–0.9**<br>12:0/12:0-PA: **0.4–2.8** | 16:0/18:1-PA: **2.5–3.7**<br>12:0/12:0-PA: **1.6–11.5** |
| LPA | 3–30 | EC 3.1.3.106 | [67–69] | 20:4-LPA: **0.2–2.2** | 20:4-LPA: **0.2–2.3** |
| C1P | 1 | EC 3.1.3.115 | [70] | d18:1/8:0-C1P: **0.08** | d18:1/8:0-C1P: **0.07** |
| S1P | *0.02–1 | EC 3.1.3.114 | [71] | d16:1-S1P: **0.010–0.48**<br>d17:1-S1P: **0.021–1.07**<br>d18:1-S1P: **0.003–0.15**<br>d20:1-S1P: **0.005–0.23** | d16:1-S1P: **0.007–0.34**<br>d17:1-S1P: **0.015–0.77**<br>d18:1-S1P: **0.002–0.11**<br>d20:1-S1P: **0.004–0.17** |
| G3P | 200–2,000 | EC 3.1.3.21 | [72, 73] | 0.7–3.7 | 0.3–1.7 |

\*Based on the report by Milo [58], the values were extrapolated as follows: protein concentration of mammalian cell, 64 g protein/L; cell volume, 2,000 $\mu\text{m}^3$ ( $2 \times 10^{-12}$ L); number of proteins per cell, $2.0 \times 10^9$ ( $3.3 \times 10^{-15}$ mol/cell); average protein length, 350 aa/protein; average amino acid molecular weight, 110 Daltons. Intracellular PC concentration and PC:PA ratio in cells are extrapolated to 20 nmol/ $10^6$ cells [59] and 50:1 [53], respectively. Total proteins in tissues (mg protein/g tissue) were calculated based on a report by Zaia *et al.* [60]. Enzyme activities of PHOSPHO2 under estimated intracellular substrate concentrations were calculated using the results (Figs. 4 and 7). Using the results, the estimated enzyme activities of PHOSPHO2 on the cellular membrane (PC-rich environment) were calculated from estimated enzyme activity in cytosol.

